# Transcriptional signatures of participant-derived neural progenitor cells and neurons implicate altered Wnt signaling in Phelan McDermid syndrome and autism

**DOI:** 10.1101/855163

**Authors:** Michael S. Breen, Andrew Browne, Gabriel E. Hoffman, Sofia Stathopoulos, Kristen Brennand, Joseph D. Buxbaum, Elodie Drapeau

## Abstract

**Background:** Phelan-McDermid syndrome (PMS) is a rare genetic disorder with high risk of autism spectrum disorder (ASD), intellectual disability and language delay, and is caused by 22q13.3 deletions or mutations in the *SHANK3* gene. To date, the molecular and pathway changes resulting from *SHANK3* haploinsufficiency in PMS remain poorly understood. Uncovering these mechanisms is critical for understanding pathobiology of PMS and, ultimately, for the development of new therapeutic interventions.

**Methods:** We developed human induced pluripotent stem cell (hiPSC)-based models of PMS by reprogramming peripheral blood samples from individuals with PMS (*n*=7) and their unaffected siblings (*n*=6). For each participant, up to three hiPSC clones were generated and differentiated into induced neural progenitor cells (iNPCs; *n*=32) and induced forebrain neurons (iNeurons; *n*=42). Genome-wide RNA-sequencing was applied to explore transcriptional differences between PMS probands and unaffected siblings.

**Results:** Transcriptome analyses identified 391 differentially expressed genes (DEGs) in iNPCs and 82 DEGs in iNeurons, when comparing cells from PMS probands and unaffected siblings (FDR <5%). Genes under-expressed in PMS were implicated in Wnt signaling, embryonic development and protein translation, while over-expressed genes were enriched for pre- and post-synaptic density genes, regulation of synaptic plasticity, and G-protein-gated potassium channel activity. Gene co-expression network analysis identified two modules in iNeurons that were over-expressed in PMS, implicating postsynaptic signaling and GDP binding, and both modules harbored a significant enrichment of genetic risk loci for developmental delay and intellectual disability. Finally, PMS-associated genes were integrated with other ASD iPSC transcriptome findings and several points of convergence were identified, indicating altered Wnt signaling, extracellular matrix and glutamatergic synapses.

**Limitations:** Given the rarity of the condition, we could not carry out experimental validation in independent biological samples. In addition, functional and morphological phenotypes caused by loss of *SHANK3* were not characterized here.

**Conclusions:** This is the largest human neural sample analyzed in PMS. Genome-wide RNA-sequencing in hiPSC-derived neural cells from individuals with PMS revealed both shared and distinct transcriptional signatures across iNPCs and iNeurons, including many genes implicated in risk for ASD, as well as specific neurobiological pathways, including the Wnt pathway.

## INTRODUCTION

Phelan-McDermid syndrome (PMS) is one of the most penetrant and more common single-locus causes of ASD, accounting for ca. 1% of ASD diagnoses [1–3]. PMS is caused by heterozygous 22q13.3 deletions or mutations leading to haploinsufficiency of the *SHANK3* gene [2, 4–6]. Clinical manifestations of PMS include ASD, global developmental delay, severe to profound intellectual disability (ID), motor abnormalities, delayed or absent speech, and epilepsy [2, 6, 7]. *SHANK3* is a scaffolding protein of the post-synaptic density of excitatory synapses and plays a critical role in synaptic function, being a key component in the integration of glutamatergic synaptic signaling [8–14]. A fundamental knowledge gap separates the well-defined clinical impact of *SHANK3* mutations on neurodevelopmental phenotypes and the molecular and cellular mechanisms leading to these phenotypes. Uncovering these mechanisms is critical for identifying drug targets and developing novel intervention strategies for PMS and for subsets of ASD that share related pathobiological mechanisms.

Several studies have utilized murine models to explore the molecular consequences of *SHANK3*-deficiency. We and others have shown that mice with a disruption in *Shank3* have altered glutamatergic signaling and synaptic dysfunction, as well as altered motor, social and repetitive behaviors [11, 15–26]. Studies on a genetically modified Shank3 rat model showed deficits in attention and in long-term social memory, which were attributable to reduced synaptic plasticity in the hippocampal-medial prefrontal cortex pathway [27, 28]. Many of the features of these rodent models reflect deficits similar to those observed in PMS. However, although animals and humans share homologous genes, pathways, and networks, rodents may have limits as models of human neurodevelopment. Specifically, the biological context and integration of molecular pathways differ across species, which can pose an obstacle for drug development and discovery.

The generation of neuronal cultures from human induced pluripotent stem cells (hiPSCs) has the potential to create translatable and experimentally tractable human neuronal models [29, 30]. hiPSCs can be derived directly from participant cells and reprogrammed to differentiate into target cell types of interest, recapitulating the early stages of neurodevelopment *in vitro*, all while retaining the genetics of the original donor. In several studies, hiPSC-derived neurons have been examined from individuals with PMS and show a reduction in the number of synapses in *SHANK3*-deficient neurons, together with impaired dendritic arborization and major deficits in excitatory, but not inhibitory, synaptic activity [31–36]. Isogenic comparisons of CRISPR-engineered heterozygous and homozygous *SHANK3* mutations demonstrated that *SHANK3*-deficiency causes functionally impaired hyperpolarization-activated cation currents, likely through its ability to interact with and organize the hyperpolarization-activated cyclic nucleotide-gated channels that mediate I_h_ currents [37]. Some studies indicate that excitatory synaptic transmission in PMS neurons can be corrected by restoring *SHANK3* expression, by treating neurons with IGF-1, or by pharmacologically and genetically activating Akt or inhibiting the Cdc2-like kinase 2 activity [31, 34, 35]. Amelioration of deficits associated with *SHANK3* haploinsufficiency have also been demonstrated by treating iPSCs with lithium or valproic acid [34].

Overall, the rodent and hiPSC-based studies consistently confirm that, at the neurophysiological level, PMS leads to a disruption in glutamatergic signaling. A next step would be to identify consistent molecular changes in PMS, specifically, the repertoire of genes and molecular pathways that are altered in expression as a consequence of *SHANK3*-deficiency. A better understanding of these molecular mechanisms may inform the search for approaches to ameliorate neurobiological and neurophysiological deficits in vitro, with the ultimate goal of advancing treatments for PMS.

The overarching objective of the current study was to identify the transcriptional signatures of *SHANK3*-deficiency in iNPCs and iNeurons by comparing genome-wide RNA-seq gene expression between PMS probands (*n*=7) and unaffected siblings (*n*=6). A multi-step analytic approach was applied to: (1) confirm the developmental specificity of our hiPSC neuronal cells; (2) quantify the variance in iNPC and iNeuron transcriptome data that is explained by differences in neural cell types, individual donors and other relevant factors; and (3) identify and characterize candidate genes, molecular pathways and co-regulatory networks associated with PMS in iNPCs and iNeurons. We identify important molecular pathways that both inform pathobiological mechanisms in PMS and suggest approaches for interventions.

## MATERIALS AND METHODS

### Participants

The study includes 13 participants (Table 1, 7 probands and 6 unaffected siblings) enrolled at the Seaver Autism Center for Research and Treatment at the Icahn School of Medicine at Mount Sinai. Individuals were referred through the Phelan-McDermid Syndrome Foundation, ongoing research studies, and communication between families. The study was approved by the Program for the Protection of Human Subjects at the Icahn School of Medicine at Mount Sinai. Parents or legal guardians provided informed consent for participation and publication.

**Table 1.**
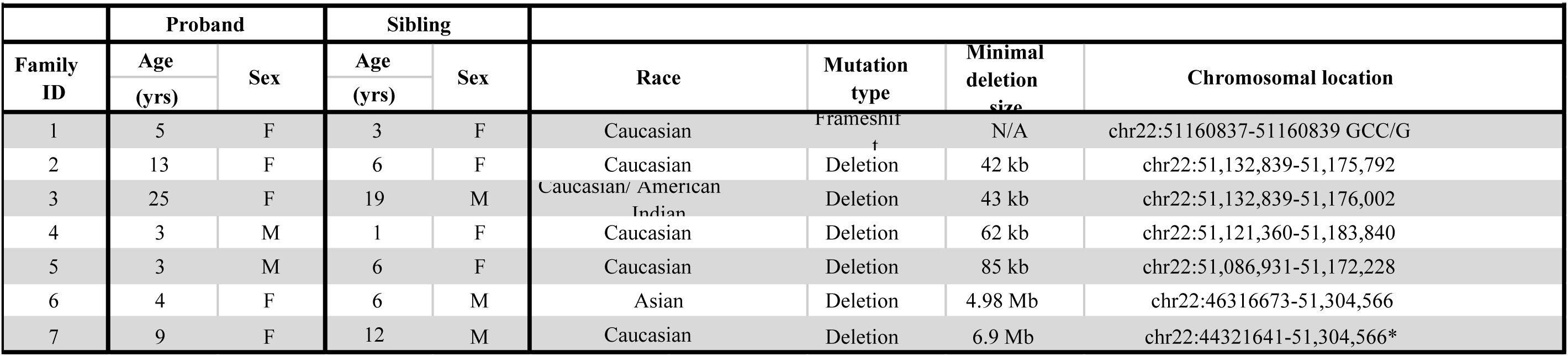
Genetic and demographic information

### Genetic findings

The mutation in patient 1 was identified through clinical WES by the Medical Genetics Laboratory at the Baylor College of Medicine. Deletions in patients 2-7 were identified as follows: Patient 2, FISH and chromosome microarray (CMA) by Signature Genomics; patient 3, CMA by the Genetics Laboratory at the University of Oklahoma Health Sciences Center; patient 4, FISH by Quest Diagnostic and CMA by the Shaare Zedek Medical Center, Jerusalem; patient 5, CMA at the UCSF Benioff Children’s Hospital Oakland; patient 6, CMA by the Mount Sinai Genetic Testing Laboratory; and, patient 7, karyotyping and custom OGT 22q array by cytogenic laboratory of the Greenwood Genetic Center.

Variants were annotated according to the Human Genome Variation Society guidelines. As reported previously, the human genome reference assembly (GRCh37/hg19 and GRCh38/hg38) is missing the beginning of exon 11 (NM_033517.1:c.1305_1346, 5′-cccgagcgggcccggcggccccggccccgcgcccggccccgg-3′, coding for 436-PSGPGGPGPAPGPG-449). We numbered nucleotide and amino acid positions according to the *SHANK3* RefSeq mRNA (NM_033517.1) and protein (NP_277052.1) sequence, in which this mistake has been corrected. Variants were interpreted according to the American College of Medical Genetics and Genomics (ACMG) guidelines.

### iPSC generation

Blood samples were collected from all participants and used for both DNA isolation (DNeasy Blood and Tissue Kit, Qiagen) and peripheral blood mononuclear cells (PBMCs) extraction (BD Vacutainer CPT Mononuclear Cell Preparation Tubes with Sodium Heparin, BD Biosciences) according to manufacturer’s instructions. PBMCs were cultured for 9 to 12 days in an erythroblast enrichment medium [38] to expand the erythroblast population and 2.5×10^5^ cells were transduced using recombinant Sendai viral vectors (Cytotune-iPSC 2.0^TM^, Thermofisher scientific), expressing the four reprogramming factors Oct4, Sox2, Kfl4 and c-Myc, according to manufacturer’s instructions. After three days, transduced cells were plated on irradiated mouse embryonic fibroblast (MEFs) and grown for two to three weeks in hiPSC medium until the emergence of individual colonies. Live hiPSCs were labeled by Tra-1-60 immunostaining (R&D systems) and positive clones were manually picked and grown on MEFs using iPSC medium. After reaching passage 10, hiPSC colonies were transitioned to feeder-free conditions using Matrigel-coated plates (Corning) and mTeSR1 medium (Stem Cell Technology) and up to three clones per individual were validated, expanded and cryopreserved.

### iPSC validation

All hiPSC lines were assessed for chromosomal abnormalities by performing karyotyping (WiCell). Their identity was confirmed using short-tandem repeat (STR) analysis (WiCell) and comparison with the donor’s blood DNA. The potential for self-renewal and pluripotency of the hiPSC lines was assessed by utilizing hiPSC RNA with the Taqman hPSC Scorecard Assay (Thermo Fisher, A15870). For pluripotency, RNA was isolated after random differentiation of iPSCs into embryoid bodies [39] to generate the three primary germ layers. Full elimination of the Sendai virus vectors was confirmed by immunostaining and with the Taqman hPSC Scorecard Assay. The e-Myco Mycoplasma PCR Detection Kit (Bulldog Bio, 25233) and the MycoAlert Mycoplasma Detection Kit (Lonza, LT07-118) were used to ensure that all the cells used in this study were mycoplasma-free.

### Generation of neuronal progenitor cells (iNPCs)

iNPCs were induced from passage 16 to P18 hiPSCs using the PSC Neural Induction Medium (Invitrogen) according to the manufacturer’s protocol. They were maintained in PSC Neural Expansion Medium up to passage 4 and then transferred to NPC medium (DMEM/F12, N2, B27 without retinoic acid, 1µg/mL natural mouse laminin, 20ng/mL FGF2). At passage 5, NPCs were labelled with Sox2 (Santa Cruz SC-17320, 1:100) and Nestin (ThermoFisher MA1-110, 1:200) antibodies to confirm their cellular identity and validated NPCs were cryopreserved. For RNA isolation, NPCs at passage 6 were seeded into 12-well plates at a density of 750,000 cells per well and harvested on the 7^th^ day after plating using RNABee (BioConnect, CS-104B) and RNA was extracted according to manufacturer’s protocol.

### Generation of forebrain neurons

For neuronal differentiation, NPCs at passage 6 were plated on to Matrigel-coated 6-well plates at a density of 200,000 cells per well in neural differentiation medium (DMEM/F12, N2, B27 without retinoic acid, 1µg/mL natural mouse laminin, 500µg/mL Dibutyryl cyclic-AMP, 20 ng/mL BDNF, 20ng/mL GDNF, 200nM L-Ascorbic Acid; [40]). Neurons were cultured for 4, 6 or 8 weeks with replacement of two thirds of the medium every 3 days. For immunostaining, additional neurons were plated on Matrigel-coated coverslips and similarly processed. At each neuronal time-point, immunohistostaining with MAP2 (Millipore MAB3418, 1:500) and Beta-3-Tubulin (Abcam ab18207, 1:1000) was performed for all samples to confirm their neuronal identity and cells for RNA sequencing were harvested using RNABee. RNA was extracted using the same procedures as described for iNPC samples. Here, the term “iNeurons” refers to mixed forebrain neuron cultures.

### RNA isolation, library preparation and sequencing

RNA samples were processed for RNA-sequencing to form two groups: 1) a larger discovery set; and 2) a smaller replication set. For the discovery set, 39 iNPC and 42 iNeuron RNA samples underwent RNA-sequencing. For the replication set, 21 iNeuron RNA samples collected at 6 weeks underwent RNA-sequencing. The integrity for each RNA sample was measured using the Agilent 2100 Bioanalyzer (Agilent, Santa Clara, CA, USA). All RNA integrity numbers (RINs) were greater than 8 (RIN: 9.59 ± 0.43). RNA samples were purified using PolyA selection, and the Illumina TruSeq Stranded Total RNA kit (Ilumina, San Diego, CA, USA) was used for library preparation, according to the manufacturer instructions. All indexed RNA libraries were pooled and sequenced using long read paired-end chemistry (2 × 150 bp) at an average read depth of ∼50M reads per sample using the Illumina HiSeq2500. Resulting short reads with Illumina adapters were trimmed and low-quality reads were filtered using TrimGalore (*--illumina* option) [41]. All high-quality reads were then processed for alignment using the hg38 reference and the ultrafast universal RNA-seq aligner STAR (v2.5.1) [42] with default parameters. Mapped bam files were sorted using Samtools and short read data were quantified using featureCounts [43] with the following parameters: -T 5, -t exon, -g gene_id. Subsequently, all read counts were exported and all downstream analyses were performed in the R statistical computing environment.

### RNA-seq data pre-processing and quality control

Raw count data was subjected to non-specific filtering to remove low-expressed genes that did not meet the requirement of a minimum of 2 counts per million (cpm) in at least ∼40% of samples. This filtering threshold was applied to iNPC and iNeuron samples separately. All expression values were converted to log_2_ RPKM and subjected to unsupervised principal component analysis (PCA) to identify and remove outlier samples that lay outside 95% confidence intervals from the grand averages as well as samples with aberrant X-inactivation gene expression profiles. A total of 32 iNPC and 42 iNeuron RNA-seq samples from the discovery set, and a total of 17 iNeuron samples from the replication set passed into downstream analyses.

### Developmental specificity analysis

Two independent analyses were performed to confirm the developmental specificity of our iNPC and iNeuron gene expression data. First, we sought to confirm the developmental origin of our samples by integrating several RNA-seq data sets from post-mortem brain tissue and hiPSC models with our iNPC and iNeuron gene expression data using a previously described analytical approach [44]. A total of 15 independent studies were collected covering 2,716 independent samples and 11,650 genes. All expression values were converted to log_2_ RPKM and collectively normalized using quantile normalization from the *limma* R package [45]. These data, along with our iNPC and iNeuron expression data were analyzed jointly and integrated using principal component analysis (PCA). Second, we sought to confirm that highly expressed genes in our current data set are indeed preferentially prenatally biased in expression, based on BrainSpan developmental RNA-seq data. We previously applied a linear regression model to 299 neocortical BrainSpan samples ranging from 8 post-conceptual weeks to 40 years of age in order to characterize 22,141 genes as either prenatally or postnatally biased (log_2_FC > 0.1 and q < 0.05) or unbiased in expression (q > 0.05) [46]. The regression model generated a ‘prenatal effect’ (*t-*statistic) of the log_2_ fold-change of prenatal versus postnatal transcript abundance. We leveraged these summary statistics to examine the top 1000 most expressed and top 1000 least expressed genes in both iNPCs and iNeurons. Each gene set was examined to determine if the distribution of the fetal effect (*i.e.* t-statistics for each gene set) differed significantly from the entire neocortical background using a Wilcoxon signed rank test. The neocortical background was defined as genes which were simultaneously detected by RNA-seq in the current study as well as genes found to be expressed in the neocortex following quality control procedures.

### Cell type deconvolution analysis

The frequencies of neural cell types were estimated using Cibersort cell type deconvolution (https://cibersort.stanford.edu/) [47]. Cibersort relies on known cell subset specific marker genes to predict the proportions of cell types in heterogeneous bulk RNA-sequencing data. The method applies linear support vector regression, a machine learning approach that is robust compared to other methods with respect to noise, unknown mixture content and closely related cell types. As input, we used a reference panel of single-cell RNA-sequencing data from the human fetal cortex [48]. Cell specific gene signatures were curated using pre-defined cell clusters from the original publication covering four cell types: *i)* dividing intermediate progenitor cells (clusters 15-19); *ii)* excitatory neurons (clusters 21-28); *iii)* inhibitory neurons (clusters 38-46); and *iv)* mixed glial cells (clusters 4,6,9,19). Definitions for excitatory and inhibitory cell lineages in these data were defined in our previous work [46].

### Quantifying transcriptome variance explained by known factors

Following data quality control, outlier detection and developmental specificity analysis (all described above), all gene expression values were normalized using VOOM normalization (a variance-stabilization transformation method) [45], and these data were used to carry out the remainder of downstream analyses. To understand the effects of various recorded factors on gene expression patterns, linear mixed effect models were applied to decompose the transcriptome variability into discrete percentages of variability attributable to multiple biological and technical sources of variation using the R package variancePartition [49]. For each gene, the percentage of gene expression variation attributable to differences in cell types, individual as a repeated measure (*i.e.*, inter-donor effects), family effects, PMS diagnosis, RIN, age, biological sex, sequencing batch and variation in estimated cell type frequencies was computed. By properly attributing multiple sources of expression variation in this fashion, it is possible to identify and partially correct for some confounding variables in our differential gene expression analysis.

### eQTL enrichment analysis

We used our previously described approach [44] to examine the overlap between genes with eQTLs from the CommonMind Consortium and genes exceeding a variance percentage cutoffs for a particular variable of interest in the current study. In brief, varianceParition analysis was applied to assign each gene a fraction of variance explained by a specific observed factor in the current analysis. A total of 40 different variance explained cutoff thresholds were examined and the overlap between genes with values exceeding this cutoff and the 2000 genes with the smallest *p*-values from cis-eQTL analysis is evaluated. The overlap is computed for the observed data and 10,000 data sets with the variance percentages randomly permutated. At each cutoff where > 100 genes are represented, the fold enrichment is computed as the observed overlap over the permuted overlap.

### Differential gene expression analysis

Differential gene expression analyses were conducted using a moderated *t*-test from the R package limma [45]. All analyses adjusted for the possible confounding influence of biological sex, sequencing batch and RIN. Moreover, due to the repeated measures study design, where individuals are represented by multiple independent iNPC and iNeuron technical replicates, the duplicateCorrelation function was applied in the limma analysis and gene level significance values were adjusted for multiple testing using the Benjamini and Hochberg method to control the false discovery rate (FDR). Genes passing a FDR < 5% were labeled as showing significantly altered expression.

### Functional enrichment of differentially expressed genes

Functional annotation was assessed in two complementary ways. First, all differentially expressed genes (FDR <5%) were functional annotated using the ToppFun module of ToppGene Suite software [50]. We explored Gene Ontology terms related to biological processes using a one-tailed hyper-geometric tested (Benjamini–Hochberg (BH) FDR corrected) to assess the significance of the overlap. Enrichment was examined separately for over-expressed and under-expressed genes. All terms must pass an FDR corrected *p*-value and a minimum of three genes per ontology were used as filters prior to pruning ontologies to less redundant terms. Second, we applied the camera function in the R package *limma* [45] to perform a competitive gene set test and to assess whether the genes in a given set are highly or lowly ranked in terms of differential gene expression relative to genes that are not in the set. The method leverages limma’s linear model framework, taking both the design matrix and contrast matrix (if present) and accommodates the observational-level weights from *voom* in the testing procedure. After adjusting the variance of the resulting gene set test statistic by a variance inflation factor that depends on the gene-wise correlation (which is set to 0.01 by default) and the size of the set, a *p*-value is returned and adjusted for multiple testing.

### Protein-protein interaction networks

The STRING database v11.0 [51] was used to assess whether differentially expressed genes were enriched for direct protein–protein interactions (PPIs) and to identify key genes mediating the regulation of multiple targets. For these analyses, our signature query of PMS-associated genes (FDR<5%) were used as input. STRING implements a scoring scheme to report the confidence level for each direct PPI (low confidence: < 0.4; medium: 0.4–0.7; high: > 0.7). We used a combined STRING score > 0.4. Hub genes within the PPI network are defined as those with the highest degree of network connections. We further used STRING to test whether the number of observed PPIs were significantly more than expected by chance using a nontrivial random background model. For visualization, the STRING network was imported into Cytoscape [52].

### CHD8 ChIP-Seq overlap analysis

To assess whether PMS-associated genes (FDR <5%) relate to known genome-wide CHD8 binding sites, we tested our differentially expressed gene sets for enrichment with human brain-specific sequences from two independent ChIP-seq studies covering: 1) 3,281 CHD8-binding sites in the human mid-fetal brain at 16-19 post-conception weeks [53]; and 2) 6,860 CHD8-binding sites in human neural progenitor cells [54] using the intersection of signal-enriched regions detected by all three CHD8 antibodies used in the study. In order to assess overlap with these binding sites, genomic coordinates were defined as the start and end positions for each differentially expressed gene (analogous to gene length). A permutation-based approach with 1,000 random permutations was used to determine statistical significance of the overlap between genomic coordinates for differentially expressed genes with CHD8-binding sites using the R package regioneR [55].

### Weighted gene co-expression network analysis (WGCNA)

Signed co-expression networks were built separately for iNPC and iNeuron samples using WGCNA [56]. To construct a global weighted network for each cell type, a total of 15,759 post QC genes across 32 iNPC samples and 16,721 genes across 42 iNeuron samples were used. The absolute values of Pearson’s correlation coefficients were calculated for all possible gene pairs within each cell type and resulting values were transformed using a β-power (β=12 for iNPC samples; β=14 for iNeuron samples) so that the final correlation matrices followed an approximate scale-free topology. The WGCNA dynamic tree-cut algorithm was used to detect network modules (minimum module size =50; cut tree height = 0.99; deep-split = 2, merge module height = 0.20). Once network modules were identified, modules were assessed for significant associations to PMS diagnosis, as well as other biological and technical factors. In order to determine which modules, and corresponding biological processes, were most associated with PMS, we ran singular value decomposition of each module’s expression matrix and used the resulting module eigengene (ME), equivalent to the first principal component, to represent the overall expression profiles for each module. This technique is useful for reducing the number of multiple comparisons from thousands of genes to tens of modules. Gene co-expression modules that were significantly associated with PMS were subjected to functional annotation using the ToppFun module of ToppGene Suite software, as described above. Fisher’s exact tests were used to assess the overlap of co-expression modules between iNPCs and iNeurons, while controlling FDR using the BH procedure.

### Curation of autism and neurodevelopmental disorder gene sets

Two tiers of gene sets were collected to examine overlap with PMS-associated genes in the current study: (1) gene sets that implicate genetic risk for ASD and neurodevelopmental disorders (NDDs); and (2) gene sets that represent differentially expressed genes induced by knockdown (KD) or knockout (KO) of an ASD or NDD gene in iPSCs. For the gene sets that cover genetic evidence for ASD and NDDs we collected loci from: i) five lists of *de novo* variants implicated in ASD [46, 57–60]; ii) loci that implicate risk for intellectual disability (ID) [61, 62]; iii) genes implicated in developmental disorders (DD) from the DDG2P database [63]. We also included genes that are direct targets of FMRP [64]. For the gene sets from other iPSC transcriptome studies, we curated previously described differentially expressed genes caused by: *i)* shRNA KD of *SHANK3* in hiPSC-derived neurons [65]; *ii)* CRISPR/Cas9 heterozygous KO of *CHD8* in hiPSC-derived NPCs and neurons [66]; *iii)* shRNA KD of *TCF4* and *EHMT1* in hiPSC-derived NPCs [67]; *iv)* shRNA KD of *MBD5* and *SATB2* in human neural stem cells [68]; *v)* shRNA KD of *NRXN1* in human neural stem cells [69]; and *vi)* CRISPR/Cas9 heterozygous and homozygous KO of ten different ASD-related genes in iPSCs and iPSC-derived neurons [70]. Full gene lists are provided in ***Supplemental Table 4***.

### Gene overlap analyses

To compute significance of all gene-based overlaps, we used a the GeneOverlap function in R which uses a Fisher’s Exact Test (FET) and an estimated odds-ratio for all pair-wise tests. Similarly, overrepresentation of ASD and NDD genetic risk gene sets within gene co-expression modules were analyzed a FET to assess the statistical significance. When testing overlap across gene modules, tests were adjusted for multiple testing using BH procedure to control the FDR.

### Availability of data and materials

RNA-sequencing fastq files have been deposited in the Gene Expression Omnibus under accession number GSEXXXX (to be released following manuscript publication).

## RESULTS

### iNPC and iNeuron RNA-seq data generation and quality control

Peripheral blood samples were reprogrammed into hiPSCs and differentiated to generate iNPCs and iNeurons from a primary cohort of individuals with PMS (*n*=7; 2 males and 5 females) and their unaffected siblings (*n*=6; 3 males and 3 females; Table 1). The majority of the PMS probands studied here harbor subtelomeric deletions spanning 40-690 kbp, with the exception of one affected individual with a *SHANK3* point mutation. For all individuals, genome-wide RNA-sequencing was generated from iNPCs and from iNeurons at four, six and eight weeks in culture, to compare transcriptional differences between PMS probands and their unaffected siblings. For each participant, 1-to-3 clones were used for the NPC and neuronal induction yielding a total of 39 iNPC samples and 42 iNeuron samples in the discovery set (**Supplemental Table 1**). Subsequently, all gene expression data were inspected for outlier samples on the basis of abnormal gene expression profiles (*i.e.* samples beyond 95% confidence interval of grand mean), and six iNPC samples were flagged and removed (Figure S1A-B). Next, because the extent of X-inactivation in females has been reported to be a quality issue during iPSC reprogramming, we examined the expression patterns of genes on the sex chromosomes using *XIST* on chrX and six genes on chrY for all samples (Figure S1C-D). This analysis identified six female iNPC samples (five of which were already removed on the basis of outlier expression profiles) that have expression patterns intermediate between males and females, consistent with either contamination or aberrant X-inactivation, which were removed from our analysis.

### Developmental and cellular specificity of iNPCs and iNeurons

We sought to determine whether our iNPC and iNeuron transcriptome data accurately reflects early developmental gene expression profiles by integrating our RNA-seq data with other studies using iPSC neuronal cell and post-mortem brain gene expression data. A total of 15 independent studies were leveraged covering 11,650 genes and 2,719 developmentally distinct samples (*see Materials and Methods*). Following standardized data pre-processing procedures, principal component analysis (PCA) stratified all gene expression samples into a distinct developmental axis starting with early embryonic stem cells and subsequently moving into iNPCs and iNeurons, and into prenatal and postnatal postmortem brain samples (Figure S2A). Embryonic stem cells (ESCs) and iPSCs clustered separately from iNPCs and iNeurons, which in turn co-clustered with early prenatal brain samples. Notably, our iNPCs and iNeurons also co-cluster with iNPC and iNeuron samples generated from previous reports confirming their early developmental gene expression profiles. This clustering was robust to differing methodologies used for iPSC reprogramming and differentiation across multiple prior studies. We also quantified whether the genes with the highest expression in our iNPC and iNeuron data sets were predominantly prenatally biased in expression using data from the BrainSpan project, and a clear prenatal bias in expression was observed for genes with the highest levels of expression across both cell types (Figure S2B). We also observed that genes with the lowest level of expression were predominantly postnatally biased in expression, indicating that markers of later postnatal brain development are expressed at low levels in the current data sets (Figure S2C).

It is possible that PMS-associated mutations could lead to unique neural cell type composition in proband, as compared to sibling, cells. In addition, genetic background or stochastic factors may also impact cell composition. We therefore estimated proportions of neural cell types for all iNPC and iNeuron samples using a reference panel of single-cell RNA-sequencing data from the fetal human cortex [48]. We observed that our iNPC samples were largely comprised of dividing intermediate progenitor cells (∼43.4%) and excitatory neurons (∼23.1%), while iNeuron samples were estimated to be comprised predominantly of excitatory neurons (∼42.4%) and inhibitory neurons (∼24.8%) (Figure S3). Comparative analyses of the estimated cell type compositions revealed minor increases in predicted proportions of excitatory neuron (*p*=0.04) and a decrease in inhibitory neurons (*p*=0.002) in PMS probands relative to unaffected siblings (Figure S3). These *in silico* predictions suggest that differences in excitatory and inhibitory cell proportions may be impacted by loss of *SHANK3*.

### Quantifying sources of gene expression variability: clinical, technical and biological factors

Inter-donor and clonal variations have previously been reported to explain a substantial fraction of gene expression variability in iPSC-derived neural cells. Therefore, as a quality check, genome-wide concordance was evaluated between technical replicates, familial related and unrelated donors. Concordance between technical replicates was examined by either origin of the same clone and the same induction or the same clone but different induction (Figure 1 A-B). Our analysis confirmed that the strongest correlation was observed between technical replicates from the same clone and same induction followed by same clone and different induction in both iNPC and iNeuron samples. Subsequently, to test the influence of various factors on gene expression profiles, for each gene, the percentage of gene expression variation attributable to each clinical and technical factor was computed. Collectively, these variables explained ∼55% of transcriptome variation, with differences between iNPC and iNeuron samples having the largest genome-wide effect that explained a median 29.2% of the observed variation, followed by differences in donor as a repeated measure (median 5.2%) and estimated excitatory cell type proportions (median 2.2%) (Figure 1C). The remaining factors explained smaller fractions of overall transcriptome variation, including family (median <0.1%) and biological sex (median <0.1%). Expression variation due to diagnosis (i.e., PMS proband) had a detectable effect in a smaller number of genes. Notably, when iNPC and iNeuron samples were analyzed separately, other technical variables such RNA integrity values, sequencing batch and total number of weeks in culture explained very little expression variation (Figure S4). Additionally, differences in *SHANK3* deletion size had a small but distinct effect on 50 genes, which were significantly overrepresented on chromosome 22 (Figure S5A-B; iNPCs, *p*=1.3e-31; iNeurons, *p*=3.2e-18). These genes were encompassed within the largest deletion reported here, and displayed clear patterns of under-expression relative to the other PMS probands (Figure S5C).

**Figure 1.**
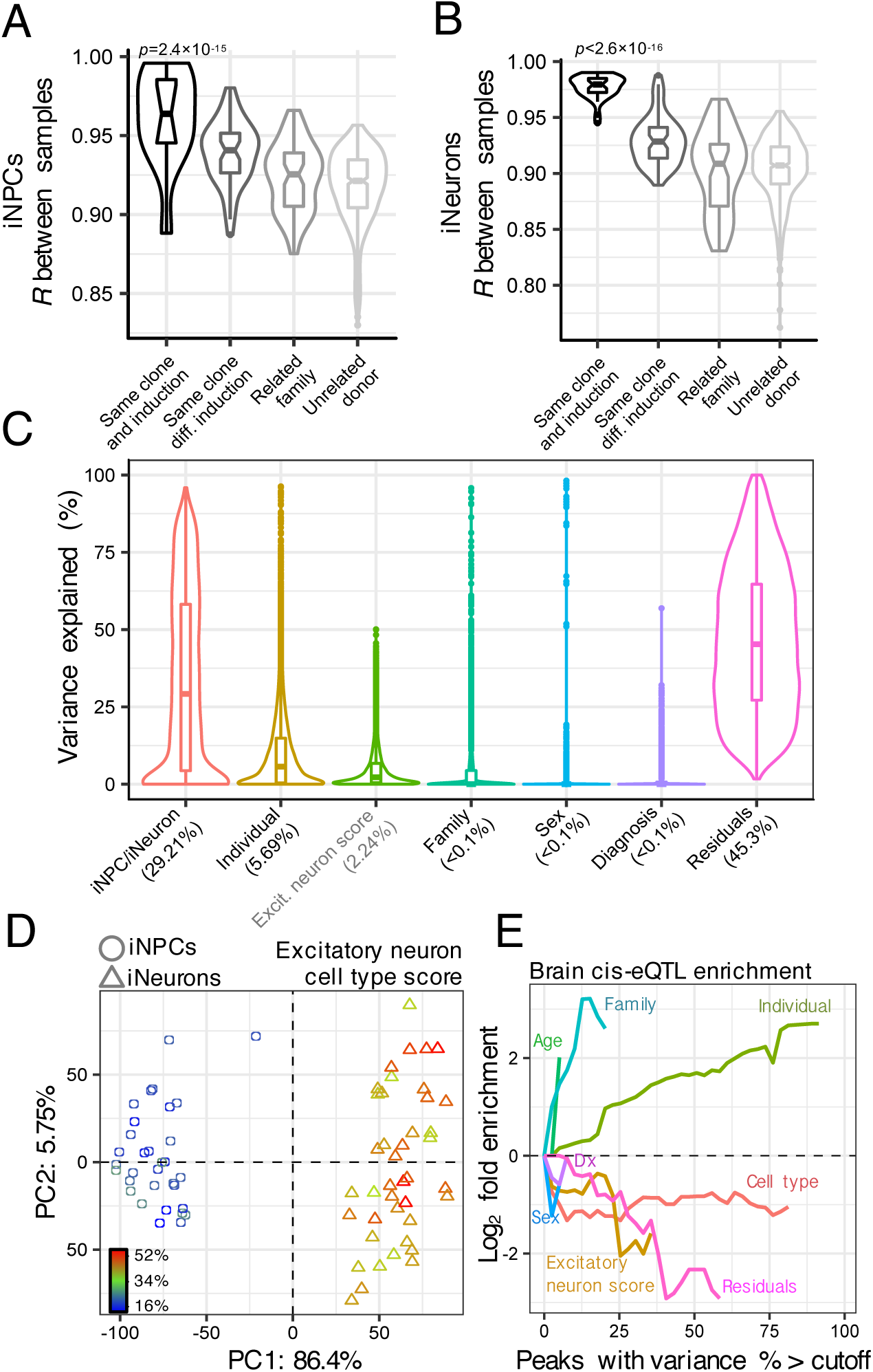
Quantifying transcriptome variance explained by observed factors. Correlation between samples from the same donor (technical replicates) compared to correlations between samples of related and unrelated family members in (**A**) iNPC and (**B**) iNeuron samples. Wilcoxon rank-sum test was used to test for differences between the means of correlation coefficients. (**C**) The linear mixed model framework of the varianceParition R package was used to compute the percentage of gene expression variance explained according to seven covariates, which represent potential biological sources of variability. Differences in cell types and excitatory neuron cell composition (estimated using CiberSort in grey) explains the largest amount of variability in the transcriptome data. (**D**) Principal components analysis (PCA) of gene expression data from iNPCs (triangles) and iNeurons (circles) where samples are colored according to their predicted excitatory neuron cell type score. (**E**) Genes that vary most across donors are enriched for brain cis-eQTLs. Fold enrichment (log_2_) for the 2000 top cis-eQTLs discovered in post mortem dorsolateral prefrontal cortex data generated by the CommonMind Consortium shown for seven sources of variation, plus residuals. Each line indicates the fold enrichment for genes with the fraction of variance explained exceeding the cutoff indicated on the *x*-axis. Enrichments are shown on the *x*-axis until less than 100 genes pass the cutoff.

Next, the influence of cell type proportions, albeit predicted, were further evaluated by overlaying excitatory neuron cell type predictions on a PCA of the gene expression data. The PCA separated iNPCs and iNeurons along the first principal component (PC), explaining 86.4% of the variance, and excitatory neuron cell estimates were separated both by PC1 and PC2 (Figure 1D). As expected, iNeuron samples had a higher proportion of predicted excitatory neurons than iNPCs (mean increase = 19.3%, *p* = 1.97e-35 by linear model), and conversely iNPCs contain a higher proportion of predicted dividing intermediate neuron progenitor cells (mean increase = 23.7%, *p* =9.16=e-63 by linear model), consistent with results derived from our previous analyses. As a final measure, we took into account the recent observation that inter-donor variation in iNPCs and iNeurons reflects genetic regulation of gene expression and shows strong enrichment for expression quantitative trait loci (eQTLs). We tested this observation in our data and similarly confirmed that genes whose variance is largely explained by differences in donor are strongly enriched for eQTLs derived from post-mortem human brain samples (Figure 1E). Variation induced by differences in iNPC and iNeuron cells and predicted cell type proportions did not reflect such genetic differences between individuals and it is likely that either stochastic or epigenetic regulators could contribute to their variability.

### Transcriptional signatures of PMS in iPSC-derived neural cells

Differential gene expression analyses comparing PMS probands and unaffected siblings identified 392 differentially expressed genes (DEGs) in iNPCs and 82 genes in iNeurons (FDR<5%; Figure 2 A-B, **Supplemental Table 2**), while adjusting for the possible influence of donor as a repeated measure, sex, RIN and sequencing batch. Genome-wide concordance was examined between iNPCs and iNeurons using PMS-associated log_2_ fold-changes, and a remarkably similar patterns of differential gene expression were observed between PMS probands and unaffected siblings in both cell types (Figure 2C; *R*=0.43, *p* <2.2e-16). Moreover, nine statistically significant DEGs were detected across both iNPCs and iNeurons, and each displayed the same direction of effect in PMS, including three genes which were consistently over-expressed (*ARHGAP20*, *PCYT2*, *CAMK2N1*) and six genes which were consistently under-expressed in PMS (*SHANK3*, *PSMD5-AS1*, *GPC3*, *TSHZ2*, *RP11-655M14.13 (lincRNA)*, *RP11-115D19.1 (lcRNA)*). Functional annotation of DEGs revealed strong pathway and biological enrichment for genes that were predominantly under-expressed in PMS in both iNPCs and iNeurons covering several early developmental terms and pathways, including 20 genes mapping to the Wnt signaling pathway (*e.g. FRZB, G3BP1, GPC3, GPC6, MLLT3, ROR2, RSPO3, WNT3A, WNT4*) (Figure 2D). Several biological processes were uniquely enriched among the under-expressed genes in iNeurons, including extracellular matrix (ECM)-related process, protein translation-related terms (*e.g,.* peptide chain elongation, translational termination/elongation/regulation/initiation) and nonsense mediated decay (Figure 2E). Overexpressed genes in iNeurons also displayed enrichment for genes involved in pre- and post-synaptic activity, cholesterol biosynthesis, transmission across chemical synapses, GABAergic synapses, G protein gated-potassium channels, signaling by insulin receptor, signaling to ERKs, glutamate binding and activation of AMPA receptors (Figure 2F). No enrichment was observed for over-expressed genes in PMS iNPC samples. Notably, adjusting for differing cell type proportions within iNPC and iNeuron samples had little effect on the resulting differential gene expression signatures (Figure S6; **Table S2**). A full table of enrichment terms can be found in ***Supplemental Table 3***.

**Figure 2.**
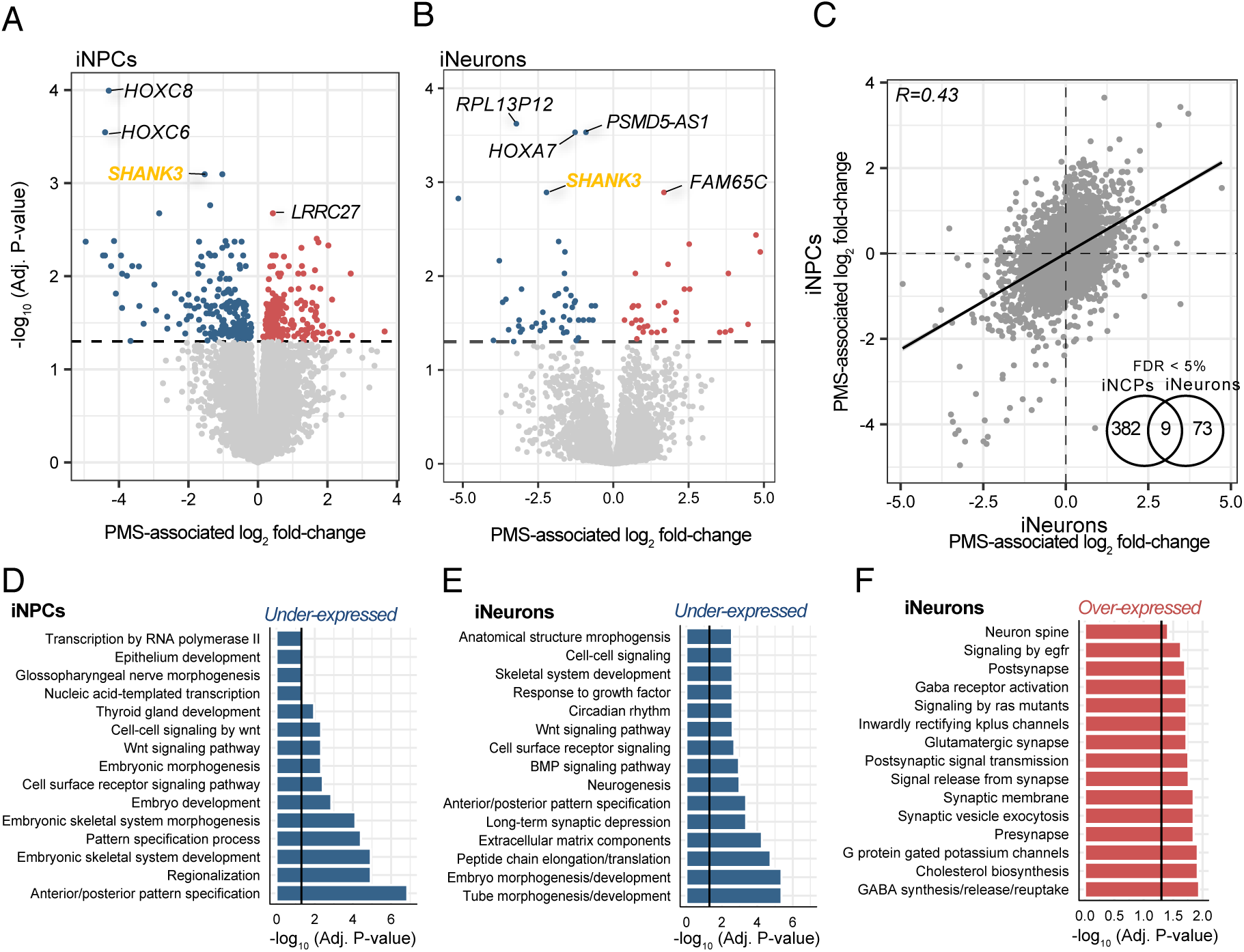
Genes and pathways associated with PMS. Differential gene expression analyses adjusted for sequencing batch, biological sex, RIN and individual donor as a repeated measure using the dupCorrelation function in the limma R package. Volcano plots compare the extent of PMS-associated log_2_ fold-changes to −log_10_ multiple test corrected *P*-value in (**A**) iNPCs and (**B**) iNeurons. Black dotted line indicates genes passing an adjusted P<0.05. (**C**) Genome-wide concordance of PMS-associated log_2_ fold-changes was examined between iNCPs and iNeurons. Inset Venn diagram displays the overlap of significant differentially expressed genes between the two cell types. Functional enrichment analysis of PMS dysregulated genes that show (**D**) under-expression in iNPCs, (**E**) under-expression in iNeurons and (**F**) over-expression in iNeurons. All enrichment terms displayed pass a multiple test corrected *P*-value.

To support these functional enrichment observations, we tested whether candidate genes that are dysregulated together indeed interact with each other at the protein level. A significant overrepresentation of direct protein-protein interactions (PPI) was identified for differentially expressed genes in iNPCs (*p*=2.32e-09, average node degree=1.81) and iNeurons (*p*= 4.19e-09, average node degree=0.81). In iNPC, hub genes in the PPI included genes involved in glutamate receptor signaling pathway, including *GRM3*, *GRIA1*, *CAMK2A* and several homeobox genes (Figure S7A). The iNeuron PPI network was notably smaller in edges and nodes, and components of the Wnt signaling pathway emerged as candidate hub genes, including *WNT3A*, *WNT7B* and *FRZB* (Figure S7B). Given that many of these genes share similar functions and interactions, we queried whether these transcriptional signatures also shared common brain-specific regulatory mechanisms. To this end, we related DEGs in PMS to well-curated binding sites for *CHD8*, a chromodomain helicase strongly associated with ASD, using *CHD8* ChIP-sequencing data from two independent studies. Among DEGs in iNPCs, significant enrichment was observed for *CHD8* binding sites derived from the human mid-fetal brain (*p*=0.03) and for binding sites derived from human NPCs (*p*=0.001). No significant enrichment for *CHD8* binding sites was observed for DEGs in iNeurons (*p*=0.46, *p*=0.41, respectively; Figure S8).

### Co-expression modules associated with PMS

Given that the majority of the PMS-associated genes share similar functions and interactions, we tested whether these genes are also co-expressed. We applied unsupervised WGCNA separately to iNPCs and iNeurons to identify small sets of genes with similar co-expression patterns. A total of 19 co-expression modules were identified in NPCs and 22 modules were identified in iNeuron samples, and all modules were well preserved between iNPCs and iNeurons (Figure S9). Each module was assessed for overrepresentation of differentially expressed genes in PMS as well as previously reported genetic risk loci for ASD and other NDDs (Figure 3A). Genes that were differentially expressed in PMS iNPCs were significantly overrepresented in iNPC module M4 (∩=53, *p*=1.53e-41), while differentially expressed genes in iNeurons were strongly enriched across three iNeuron modules: M2 (∩=12, *p*=0.003), M4 (∩=14, *p*=2.6e-5) and M19 (∩=32, *p*=2.15e-21). Notably, module M2 in iNeurons harbored a significant fraction of genetic risk loci for ID (∩=5, *p*=0.02), while module M4 in iNeurons was enriched for DD (∩=14, *p*=0.02) and ID (∩=5, *p*=0.002) risk loci. Module eigengene (ME) values for all modules were then regressed onto individual diagnostic status (*i.e.* PMS probands), which confirmed significant module-trait associations for module M4 in iNPCs with PMS (*r*=-0.58, *p*=6e-04) as well as iNeuron modules M2 (*r*=0.50, *p*=9e-04), M4 (*r*=0.55, *p*=2e-04) and M19 (*r*=-0.55, *p*=2e-04) with PMS (Figure 3B). Next, the gene-module assignments identified for iNPC and iNeuron samples, respectively, were used to perform supervised module construction for the same set of genes in the contrasting cell type (*i.e.* genes in module M1 identified in iNPCs were forced to form a module in iNeurons), which were similarly tested for association with PMS. In doing so, we found that genes that were either negatively or positively associated with PMS in one cell type, displayed similar levels of association to PMS in the other cell type (Figure 3B), consistent with our differential gene expression analysis (Figure 2C). Functional annotation of these candidate modules revealed similar biological functions as previously reported from differential gene expression, including under-expression of iNPC module M4 and iNeuron module, which were both enriched for early embryonic development gene sets, ECM, neurogenesis and Wnt signaling (Figure 3C). In iNeurons, module M2 was positively associated with PMS and was implicated in GDP binding, response to oxygen/stress/hormones, LRR domain binding. A separate iNeuron module M4 was enriched for GTPase signaling, postsynaptic signal transduction and axon guidance-related processes.

**Figure 3.**
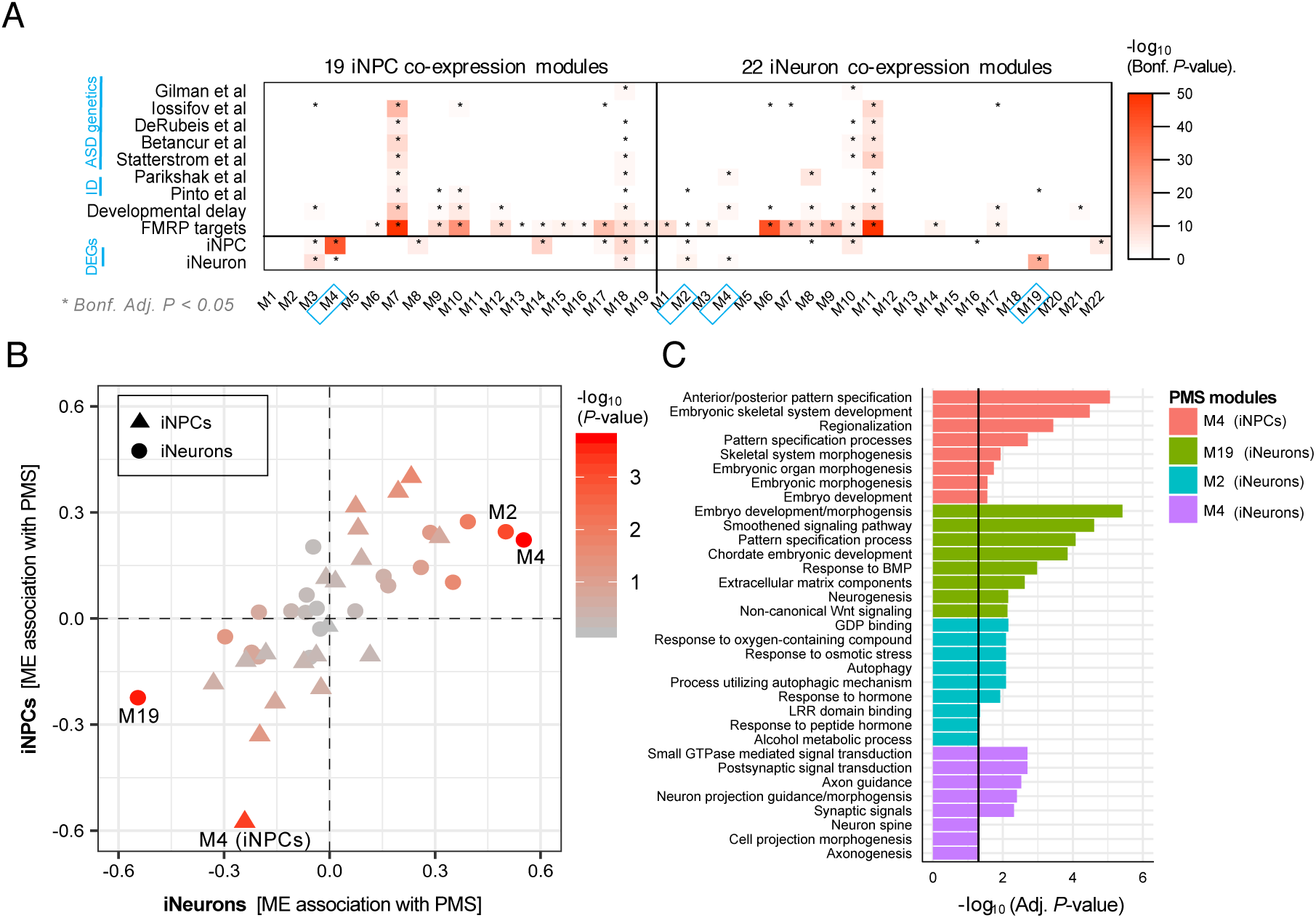
Genes co-expression analysis and enrichment. (**A**) A total of 19 co-expression modules were identified in iNPC samples and 22 modules were identified in iNeuron samples, and each module was tested for enrichment of genetic risk loci for ASD, ID and DD using findings from other large-scale studies. Modules were also examined for enrichment of target genes of FMRP, an RNA binding protein that is associated with ASD risk, as well as differentially expressed genes identified in the current study (see Figure 2). Enrichment was assessed using a Fisher’s exact test to assess the statistical significance and *p*-values were adjusted for multiple testing using the Bonferroni procedure. We required an adjusted *P*-value < 0.05 (*) to claim that a gene set is enriched within a user-defined list of genes. (**B**) Module eigengene (ME) values were associated with PMS for iNPCs (triangles) and iNeurons (circles). Next, genes in the iNPC samples were then forced to construct modules using the gene-module assignments identified in the iNeuron samples, and vice versa, and these ME values were also associated with PMS. (**C**) Functional enrichment was performed on four PMS-associated modules and the top eight enrichment terms (removing redundant annotations) are displayed.

### Overlap with existing ASD transcriptome iPSC reports

We also explored points of convergence between our *SHANK3*-deficiency findings in PMS with gene expression changes induced by either shRNA knockdown (KD) or CRISPR/Cas9 knockout (KO) of other top ranked ASD genes assayed in neural cell types (Table 2; Figure S10). A total of 6 iPSC transcriptome studies spanning 17 different ASD and NDD genes were evaluated. We identified several significant overlaps between PMS-associated gene findings in both iNPCs and iNeurons with gene expression perturbations associated with: *i) SHANK3* KD in neurons (∩=20, FET=2.4e-7; ∩=44, FET=0.003, respectively); *ii) CHD8* KO in NPCs (∩=33, FET=0.004; ∩=15, FET=0.1.8e-5, respectively); *iii) CHD8* KO in neurons (∩=91, FET=1.3e-7; ∩=29, FET=5.7e-7, respectively); *iv) EHMT1* KD in NPCs (∩=23, FET=0.001; ∩=10, FET=0.001, respectively); *v) NRXN1* KD in stem cells (∩=8, FET=0.001; ∩=4, FET=0.04, respectively); *vi) SCN2A* KO in iPSCs (∩=55, *p*=0.0002; ∩=16, *p*=0.001, respectively); *vii) ATRX* KO in iPSCs (∩=32, FET=0.03; ∩=10, FET=0.02, respectively); and *viii) ATRX* KO in neurons (∩=47, FET=6.73e-7; ∩=12, FET=0.002, respectively). In addition, differentially expressed genes in PMS iNeurons, but not in iNPCs, were enriched for genes associated with: *ix) SATB2* KD in stem cells (∩=8, FET=0.0002); and *x) TNEM1* KO in neurons (∩=7, *p*=0.004). Importantly, the vast majority of these observed overlaps were consistently enriched for genes implicating changes in Wnt signaling, ECM, perineuronal net and glutamatergic synapses (Table 2; Figure S10).

**Table 2.**
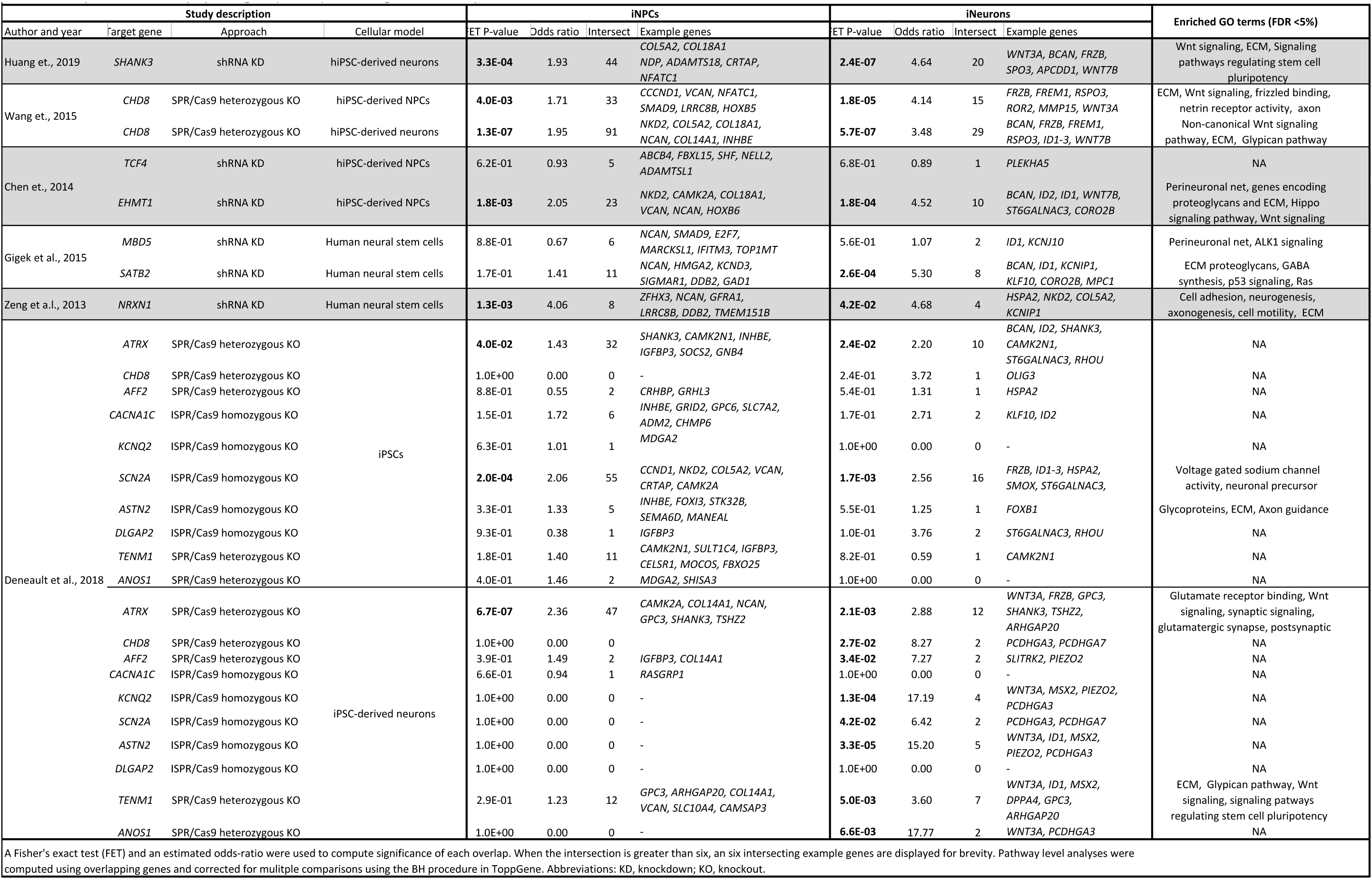
Overlap of PMS differentially expressed genes (FDR <5%) with exisiting ASD iPSC transcriptome studies.

### Validation of PMS-associated gene dysregulation in iNeurons

To validate our PMS transcriptional signatures, we performed additional RNA-sequencing on a replication set of 21 iNeurons collected at 6 weeks. Following data preprocessing, four samples were removed on the basis of aberrant X-inactivation (Figure S11A-B) and a total of eight biological replicates derived from independent differentiations and nine technical replicates passed into our subsequent validation analyses. *In silico* predictions of cell type frequencies validated the trending decreases in inhibitory neurons (*p*=0.05) and increases in excitatory neurons (*p*=0.08), albeit to an insignificant extent in PMS (Figure S11C). Subsequently, sample-to-sample correlation coefficients were evaluated between discovery and replication samples, first among technical replicates (median *R*=0.98), then among biological replicates derived from independent differentiations (median *R*=0.96) and between unrelated donors (median *R*=0.94) (Figure 4A). Overall, levels of concordance were highest between technical replicates relative to those observed between biological replicates (*p*=2.6e-9) and unrelated donors (*p*=7.7e-12). Next, differentially expressed genes were computed using the replication set of iNeurons and genome-wide concordance of PMS-associated log_2_ fold-changes were regressed onto log_2_ fold-changes computed using different combinations of discovery set iNeuron samples: i) six week samples; ii) four and eight week samples; or iii) four, six and eight week samples. As expected, the highest levels of concordance were observed between discovery and replication six week samples (*R*=0.92) followed by a combination of four, six and eight weeks (*R*=0.86) and subsequently four and eight week samples (*R*=0.80) (Figure 4B).

**Figure 4.**
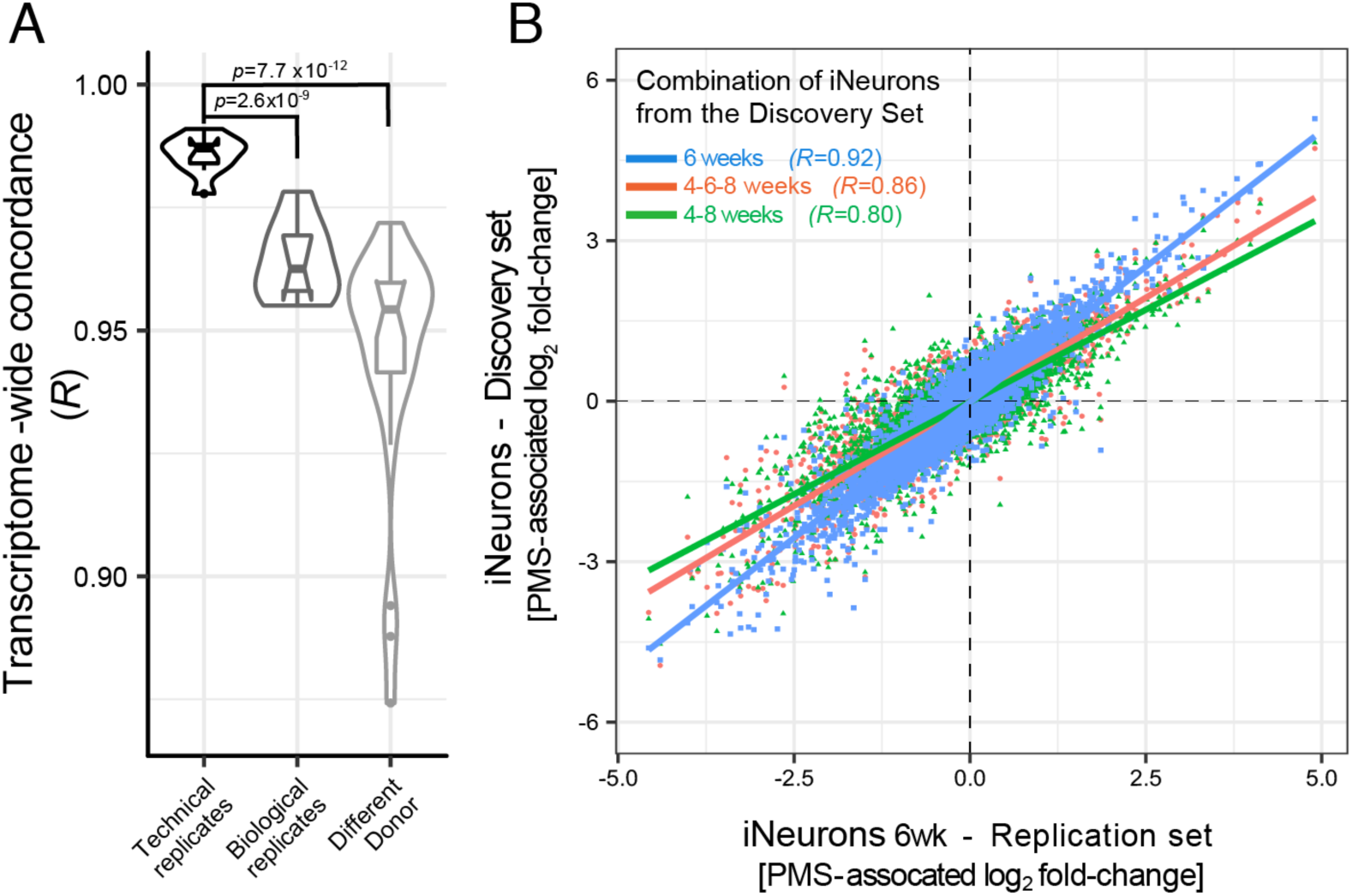
Replication of iNeuron RNA-seq. A replication set of iNeuron samples collected at 6 weeks in culture were subjected to RNA-seq. (**A**) Correlation coefficients between samples from the same donor and same clone (technical replicates), biological replicates from independent inductions and correlations between all other samples. A Wilcoxon rank-sum test was used to test for differences between the means of correlation coefficients. (**B**) The second replication batch of iNeuron samples were used to derive differential gene expression signatures between PMS probands and unaffected siblings. The PMS-associated log_2_ fold-changes from this replication set (x-axis) were compared to PMS-associated log_2_ fold-changes from the discovery set of samples, which were derived using combinations technical replicates and biological replicates at different weeks in culture (y-axis).

## DISCUSSION

Our study sought to characterize transcriptional signatures of *SHANK3* haploinsufficiency in neurodevelopment, by comparing genome-wide RNA-seq profiles of iNPCs and iNeurons derived from individuals with PMS with those of their unaffected siblings. We report on the largest sample set of PMS-derived iNPCs and iNeurons to date, representative of a range of genetic lesions associated with PMS, from a *SHANK3* point mutation to small and large 22q13.3 deletions. DEGs in our dataset were enriched for pathways involved in core developmental processes such as pattern specification and embryonic morphogenesis, including Wnt signaling pathways that are essential for neuronal fate specification. Gene co-expression modules generated from these data demonstrated convergence between altered PMS molecular pathways and ASD and NDD genetic risk loci. Importantly, overlapping DEG findings were identified between the current study and findings from other ASD and NDD transcriptome iPSC studies, demonstrating overlapping changes in RNA involved in Wnt signaling, ECM and glutamatergic synapses.

The transcriptional signatures of PMS in iNeurons point to altered postsynaptic density, glutamatergic synaptic and GABAergic genes. Our results are in line with evidence supporting a role for *SHANK3* prior to synaptogenesis and neural circuit formation, specifically in early morphogenesis and excitatory/ inhibitory balance [32, 65, 71–74]. For example, a zebrafish model of PMS that utilized morpholinos to disrupt shank3a and shank3b resulted in delayed mid- and hindbrain development, disruptions in motor behaviors, and seizure-like behaviors [73].

*SHANK3* may mediate presynaptic function via transynaptic signaling through cell adhesion molecules such as neurexin and neuroligin [75, 76]. In a rat hippocampal *in vitro* model, *SHANK3* expression was found to affect transynaptic signaling by modulating pre- and postsynaptic protein content and neurotransmission efficiency through neurexin-neuroligin interactions [74]. *SHANK3* has been shown to bind neuroligin via its PDZ domain [77], therefore potentially regulating synaptic strength via retrograde signaling through cell adhesion molecules. In addition, since neurexin-neuroligin are implicated in the regulation and coordination of synaptic function via transynaptic signaling [76], with some evidence of NMDAR regulation involvement [78], their association with *SHANK3* presents with one possible mechanism by which *SHANK3* disruption could dysregulate excitatory/inhibitory balance in the developing brain.

Our group has previously shown that *SHANK3* point mutations are sufficient to convey a PMS phenotype [4], although larger deletion sizes have been associated with a more severe range of PMS manifestations [6, 7, 41]. Here, we find that differences in *SHANK3* deletion size have a significant dosage effect on 50 genes that span the largest deletion in our dataset. One of these genes, *WNT7B*, had been previously associated with macrocephaly and chromosome 22 deletions greater than 5Mb [7]. *WNT7B* codes for a secreted signaling protein that is central to Wnt signaling pathway, which is also enriched in our dataset and has been implicated in *SHANK3* deficiency in previous reports [79]. These findings suggest that genes in close proximity to *SHANK3* on chromosome 22 may also play a role in modulating the pathobiology of PMS.

We also found several points of convergence on ECM and Wnt signaling in PMS and other iPSC studies of ASD and NDDs. Numerous lines of evidence point to Wnt signaling as a candidate pathway implicated in ASD etiology [80]. In previous reports, mutations of Wnt signaling pathway genes involved in processes such as neurite growth, synapse formation, neurogenesis and corticogenesis have been associated with ASD phenotypes [81, 82]. For example, *CTNNB1*, which produces the protein β-catenin, plays a critical role in cell adhesion and cell signaling in the Wnt signaling pathway and *de novo* mutations in *CTNNB1* have been linked to individuals with DD, ID and ASD [46]. In murine models, stabilization of *CTNNB1* in cortical samples has been found to increase Wnt signaling and boost neurogenesis [83], while depletion of *CTNNB1* from inhibitory neurons leads to deficits in neuronal activation and ASD-like behavior [84]. Numerous top-ranked ASD risk genes have also been found to function with *CTNNB1* and the Wnt signaling pathway. For example, *CHD8* is a chromatin remodeling factor and a top-ranked ASD risk gene, which has been shown to be a positive regulator of *CTNNB1*-mediated Wnt signaling in NPCs [85]. Notably, many under-expressed genes in the current dataset show enrichment for both Wnt signaling genes and *CHD8* binding sites. Both *PTEN* and *TCF7L2* also represent ASD and ID risk genes [58, 60, 86, 87], respectively, and have been identified to function with *CTNNB1* to regulate normal brain growth [88] and to initiate transcriptional responses following Wnt receptor binding [89]. Additionally, *de novo* mutations in *DDX3X* are associated with ID and ASD [90], and this gene has been recently identified to be an important component of *CTNNB1*-mediated Wnt signaling by regulating kinase activity, which in turn promote phosphorylation of Dvl and represents a major hub in the Wnt pathway [91]. Therefore, in addition to ASD, *CTNNB1*-mediated Wnt signaling may be disrupted in ID and other NDDs, further underscoring the points of convergence identified the current study and demonstrating the importance of this pathway in proper neurodevelopment.

Given the described changes in the Wnt signaling pathway, a follow-up question would be whether pharmacological regulation of Wnt signaling represents an important and/or plausible treatment strategy for PMS and ASD. Notably, several medications have been shown to modulate Wnt signaling, including methylphenidates [92], selective serotonin reuptake inhibitors (SSRIs) [93] and some antipsychotic medications [89, 94]. For example, long-term administration of methylphenidate in mice has been shown to modulate key components of the Wnt signaling pathway, including Akt and GSK3 [92]. Similarly, SSRI treatment (*e.g.* fluoxetine) has been shown to boost Wnt signaling, specifically *Wnt2* and *Wnt3* in two different mouse studies [93, 95]. Further, two additional reports have also demonstrated that the antipsychotic medication haloperidol promotes Wnt signalling, including *WNT5A* and β-catenin expression [96] as well as the phosphorylation of Akt [94]. Overall, it is noteworthy that several pharmacological interventions for behavioral disorders affect components of Wnt signaling, either directly or indirectly. However, additional investigations are required to determine the role of these mechanisms and compare the treatment efficacy across individuals with and without Wnt signaling abnormalities.

The current study also presents some limitations. First, given the rarity of PMS, we could not carry out experimental validation in independent biological samples. Nevertheless, in an effort to boost signal over noise, several points of convergence were identified with gene-based findings from other iPSC transcriptome studies. Second, loss of *SHANK3* has been shown to affect neurite length, complexity of neurite arborization and soma area, which were not examined in the current study and may contribute to some of the observed transcriptional changes. Third, while *in silico* predictions of cell type proportions attempted to control and quantify the variance in these transcriptome data, it remains possible that some transcriptional changes can be related to changes in proportions of specific cell types. This is especially true when studying iNeurons, which reflect a heterogeneous mixture of neuronal subpopulations and mixed glial cells. Single-cell RNA-sequencing of iNeurons at different differentiation stages may produce a clearer picture of the underlying cellular heterogeneity and corresponding gene expression profiles in such samples.

In summary, our study demonstrates that *SHANK3*-deficiency results in profound transcriptional changes in PMS-derived hiPSC-iNPCs and hiPSC-iNeurons. Many early developmental pathways are impacted, including altered processes related to pre-and post-synaptic signaling, embryonic development and function, as well as Wnt and ECM signaling. Several other iPSC transcriptome studies of ASD and NDD genes also displayed changes in ECM and Wnt signaling, providing molecular insights into PMS and into NDDs more broadly.

## DECLARATIONS

## Acknowledgements

We would like to thank the families who kindly consented to participate in our study.

## Authors’ contributions

JDB, AB and ED conceived of the project. AB and ED prepared the iPSC lines, neuronal differentiation and RNA-seq samples. MSB performed the computational data analysis and wrote the manuscript with SS, GH, KB, JDB and ED. All authors edited and approved the final manuscript.

## Ethics approval and consent to participate

All study participants have given written informed consent, and the genetic study has been approved by the Icahn School of Medicine at Mount Sinai Institutional Review Board.

## SUPPLEMENTAL FIGURES AND LEGENDS

**Figure S1.**
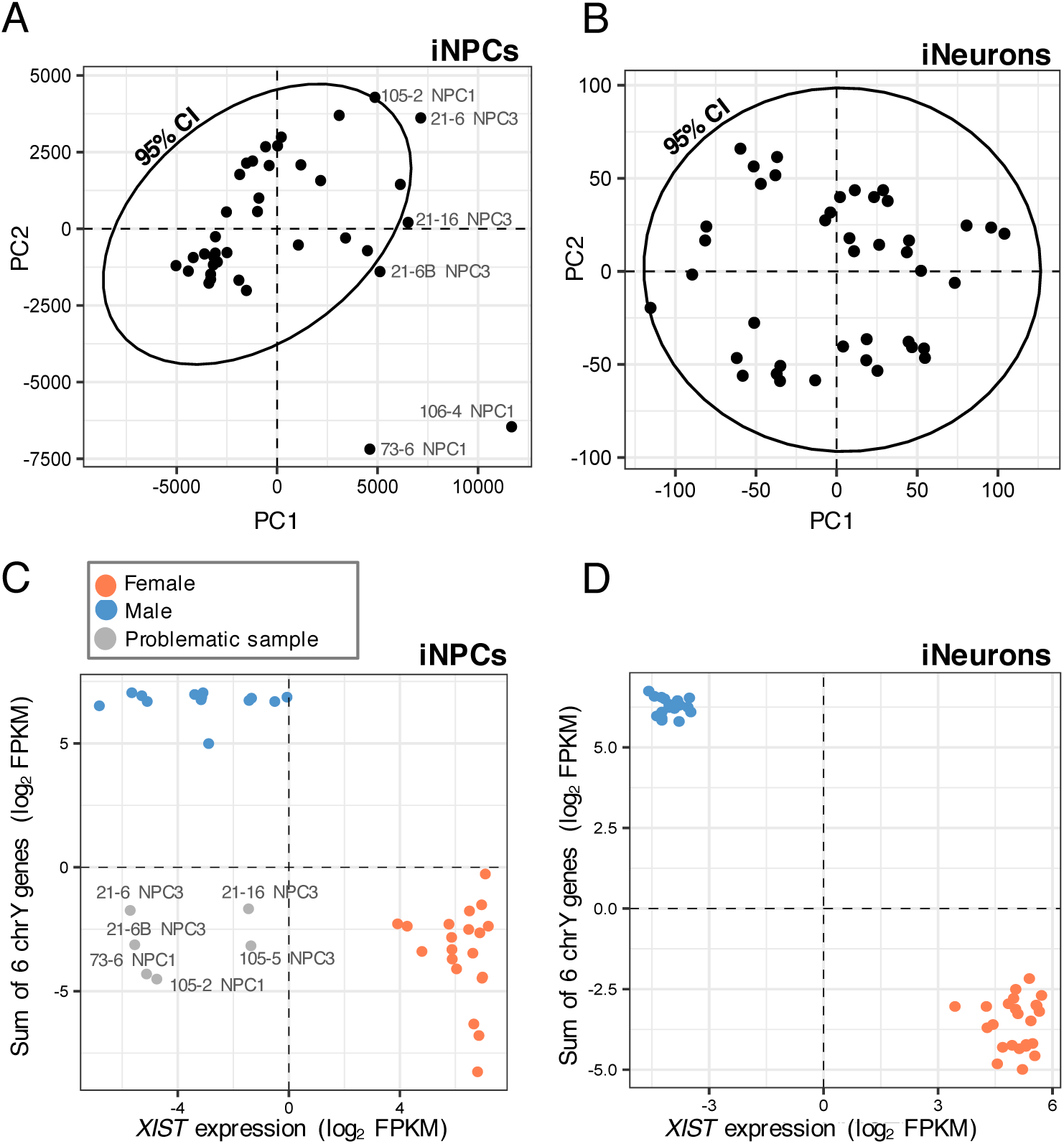
Outlier analyses. Principal component analyses were performed on RPKM values for all (**A**) iNPC and (**B**) iNeuron gene expression samples. Outliers beyond the 95% confidence intervals (black ellipse) were excluded from downstream analyses. We also sought to identify samples that may have under-gone issues with X-inactivation and/or sample mislabeling by confirming that the reported biological sex is concordant with gene expression on chrX and chrY for both (**C**) iNPC and (**D**) iNeuron samples. The expression on XIST from chrX was plotted against the sum of expression of six chrY genes (*USP9Y*, *UTY*, *NLGN4Y*, *ZFY*, *RPS4Y1*, *TXLNG2P*). Female samples with intermediate expression profiles were excluded from further analysis.

**Figure S2.**
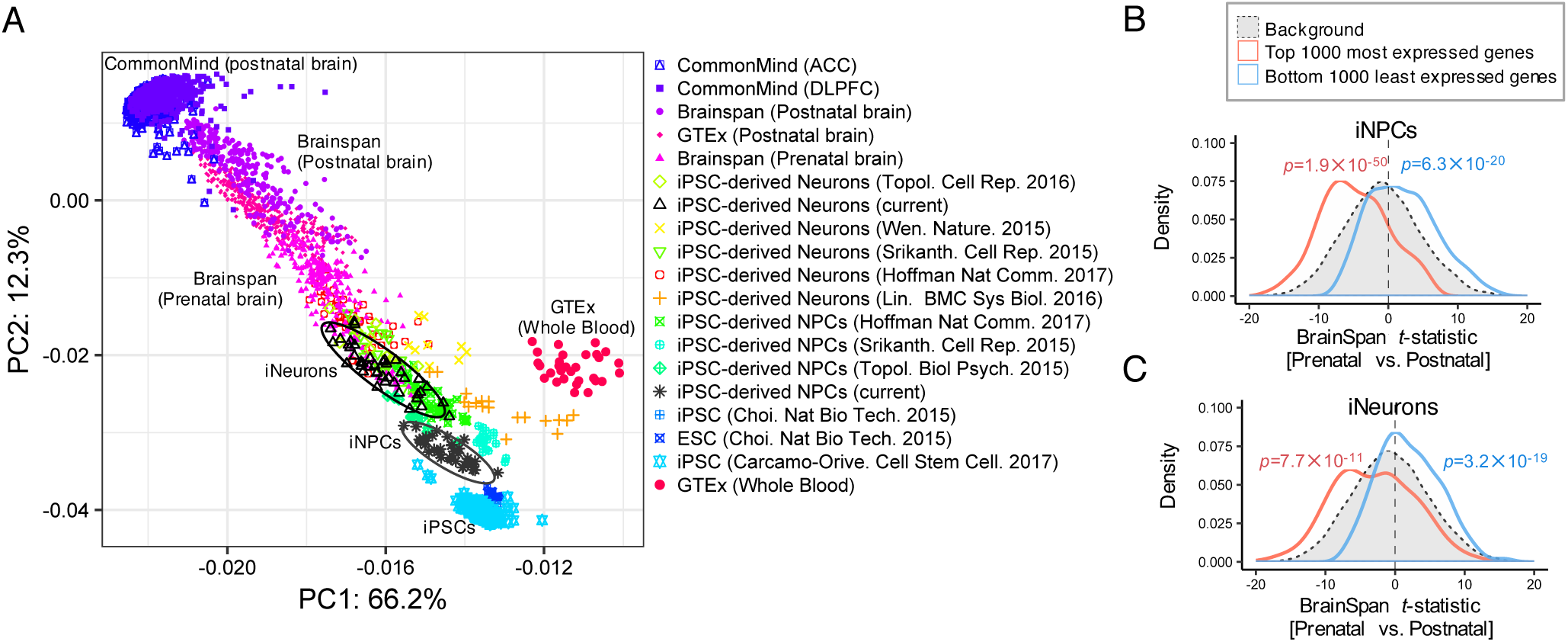
Developmental specificity analysis. (**A**) Several postmortem brain and hiPSC RNA-seq data sets spanning a broad range of developmentally distinct samples were integrated with the hiPSC-derived iNPCs and iNeurons in the current study by principal component analysis to confirm their developmental specificity. The first two principal components are shown and the iNPC samples (black stars) and iNeuron samples (black triangles) are each outlined by 95% confidence intervals. A t-statistic was calculated comparing prenatal to postnatal expression in the BrainSpan bulk RNA-seq data. (**B**) In iNPC samples, the t-statistic distribution of the top 1000 most expressed shows a prenatal bias and the top 1000 least expressed genes shows a clear postnatal bias. (**C**) A similar pattern was observed for the top 1000 most and least expressed genes across iNeuron samples.

**Figure S3.**
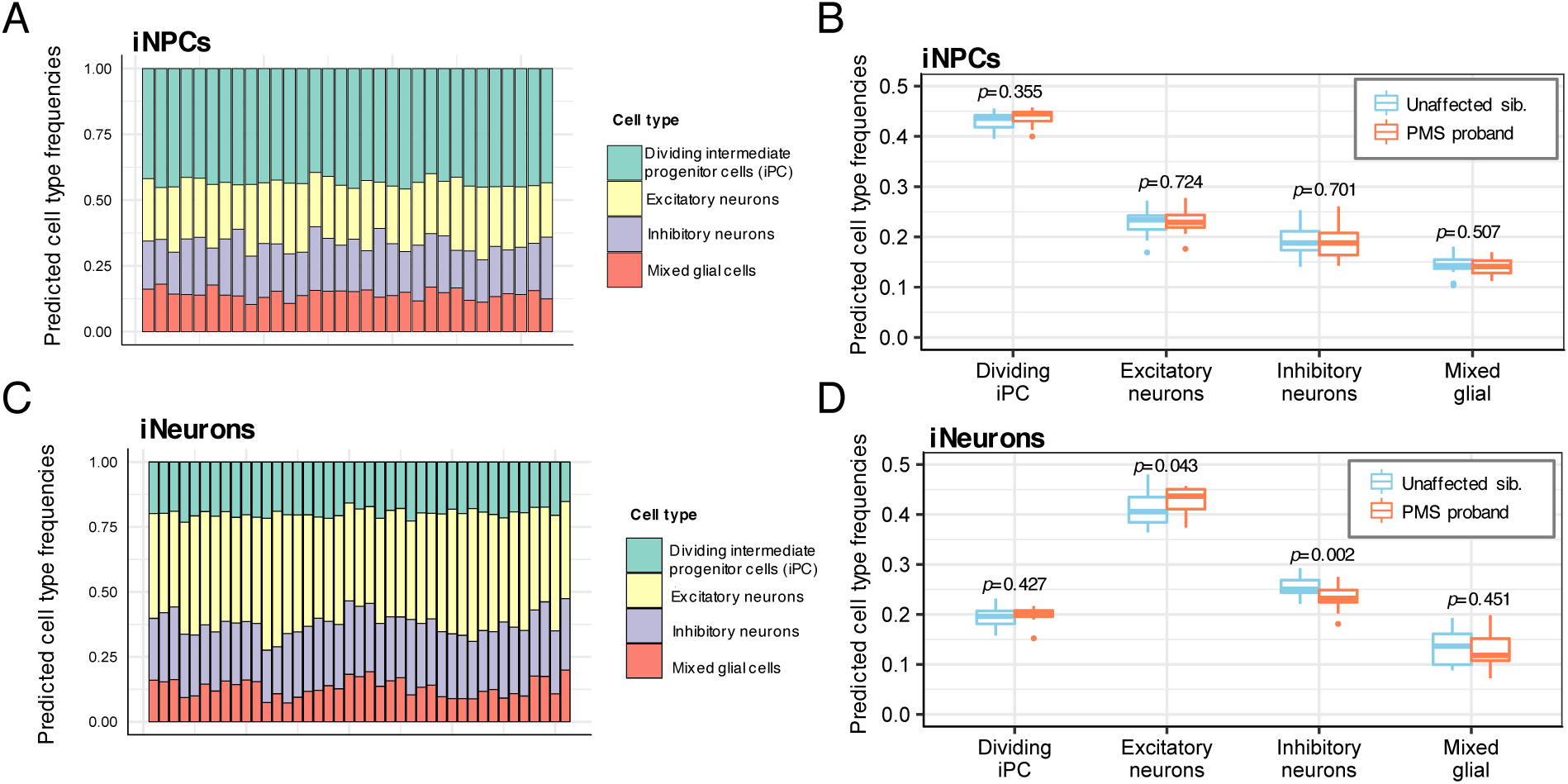
Cell type deconvolution analysis. Cibersort cell type deconvolution analysis of global gene expression profiles estimated cell frequencies (y-axis) in (**A-B**) iNPCs and (**C-D**) iNeurons for four major cell types (x-axis) using a reference panel of single-cell RNA-sequencing data from the human fetal cortex. The predicted cellular proportions were compared between PMS probands and unaffected siblings to confirm that major shifts in underlying cell types would not confound downstream analyses. A Wilcox rank-sum test was used to compare the fractions of cell proportions between probands and siblings.

**Figure S4.**
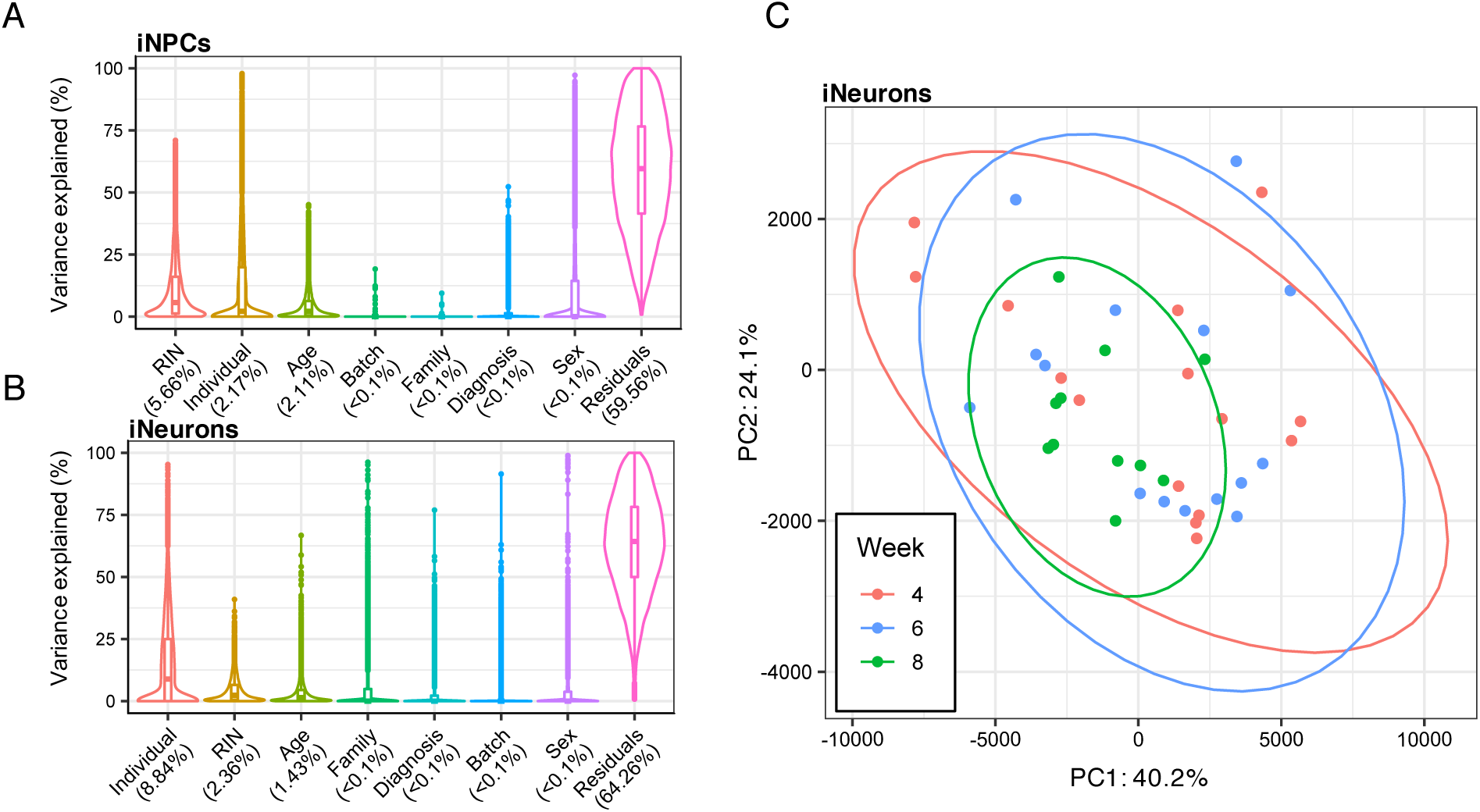
Variance explained by technical factors. The linear mixed model framework of the varianceParition R package was used to compute the percentage of gene expression variance explained by multiple biological and technical factors for (**A**) iNPCs and (**B**) iNeurons. (**C**) The variance explained by the total number of weeks iNeurons spent in culture was further evaluated by principal component analysis.

**Figure S5.**
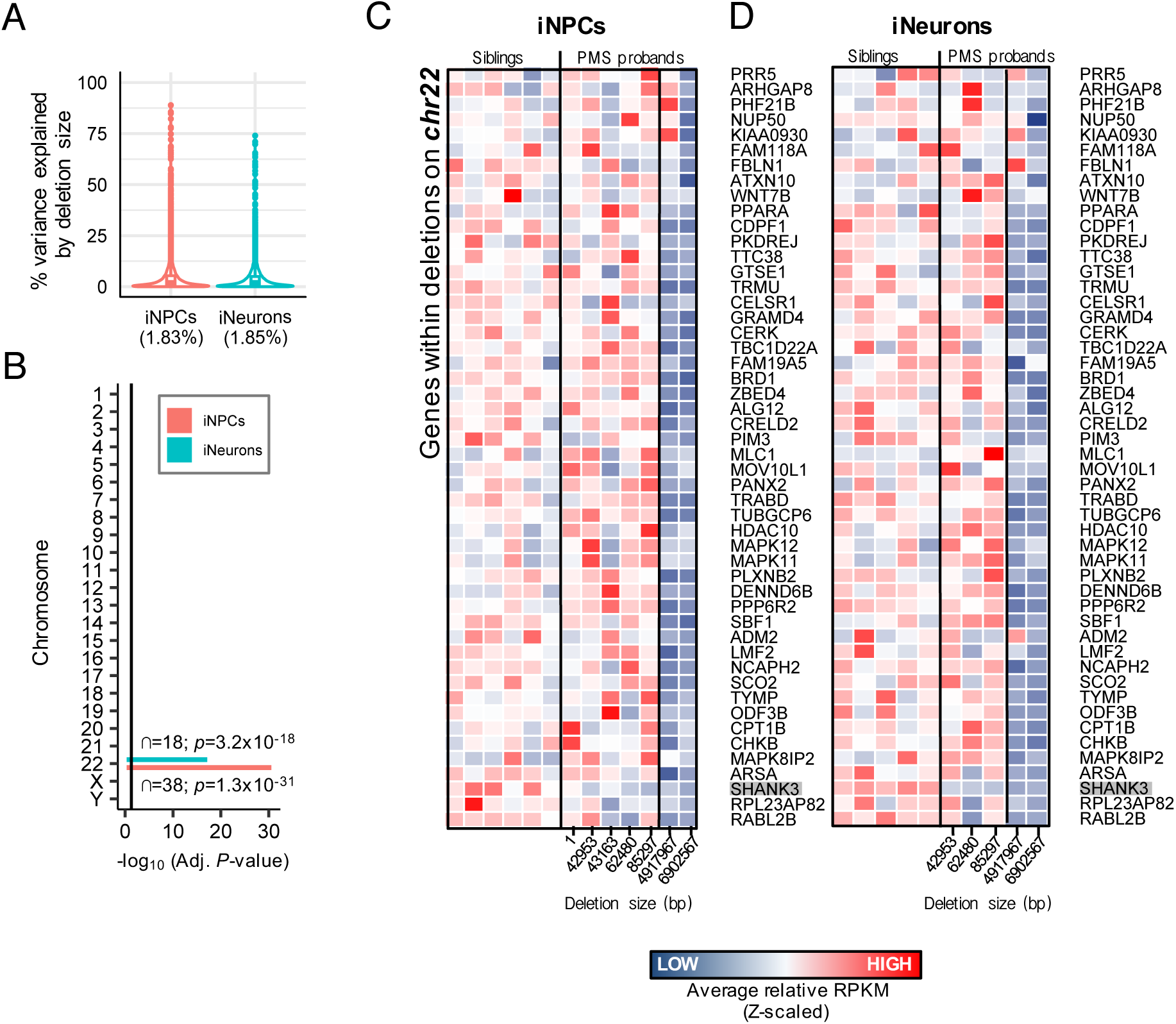
Variance explained by SHANK3 deletion size. (**A**) The linear mixed model framework of the varianceParition R package was used to compute the percentage of gene expression variance explained by SHANK3 deletion size in iNPCs and iNeurons. (**B**) Genes with variance explained >50% by deletion size were examined for chromosomal enrichment, and strong enrichment for chromosome 22 was observed. The vertical black line indicates −log_10_ P-value < 0.05. (**C**) Fifty unique genes were identified that varied by deletion size and mapped to chromosome 22, which were plotted on a heatmap using average expression values across all technical replicates for iNPC and iNeuron samples. The size of *SHANK3* deletion (bp) is displayed on the x-axis.

**Figure S6.**
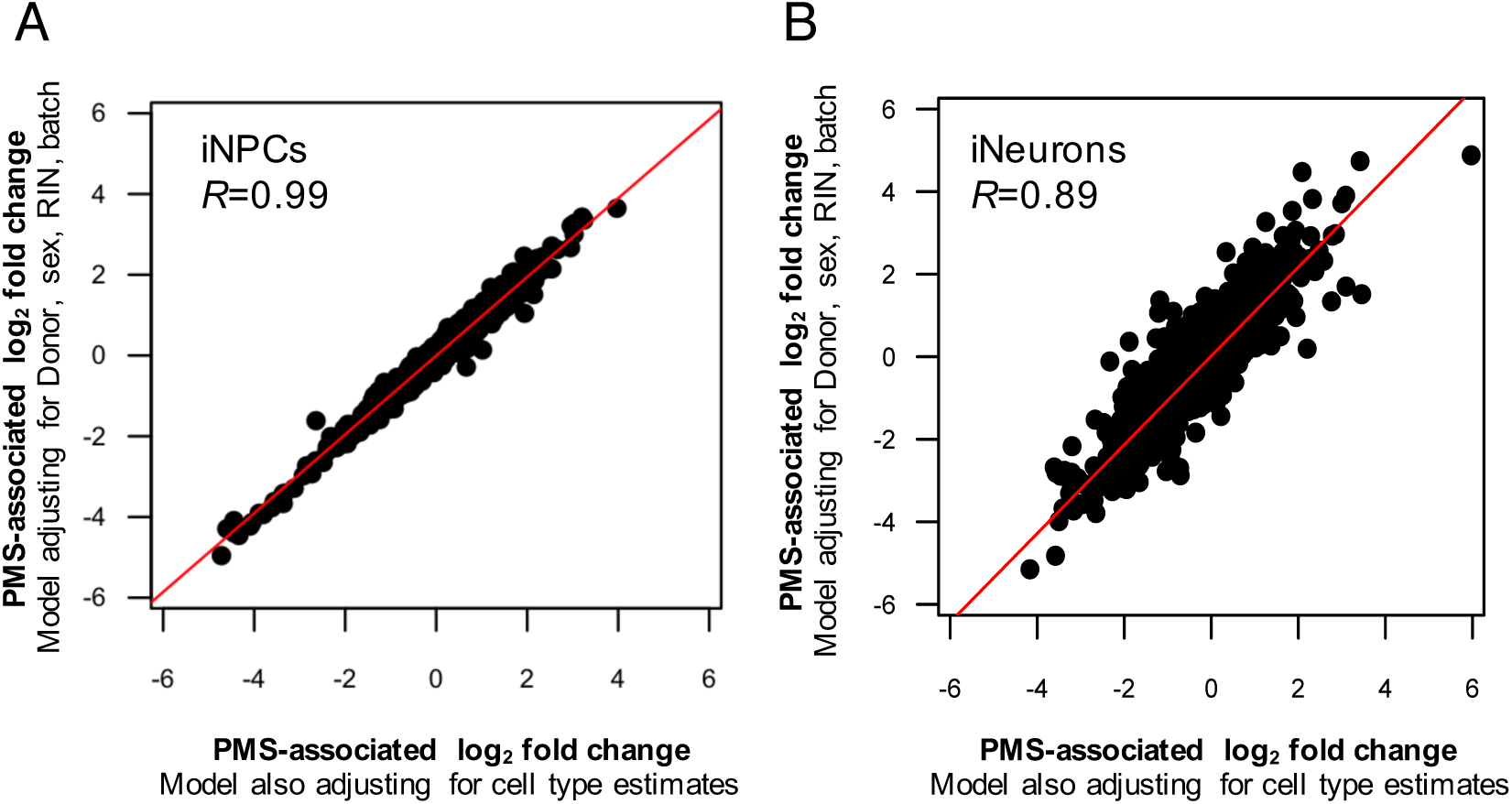
Controlling for cell type frequencies for differential expression. The concordance of genome-wide PMS-associated log_2_ fold-changes were evaluated comparing two models: i) one model adjusting for sequencing batch, biological sex, RIN and individual donor as a repeated measure on the y-axis; and ii) a second model adjusting for the same factors plus predicted excitatory neuron cell type composition on the x-axis. Concordance was examined for both (**A**) iNPC and (**B**) iNeuron samples.

**Figure S7.**
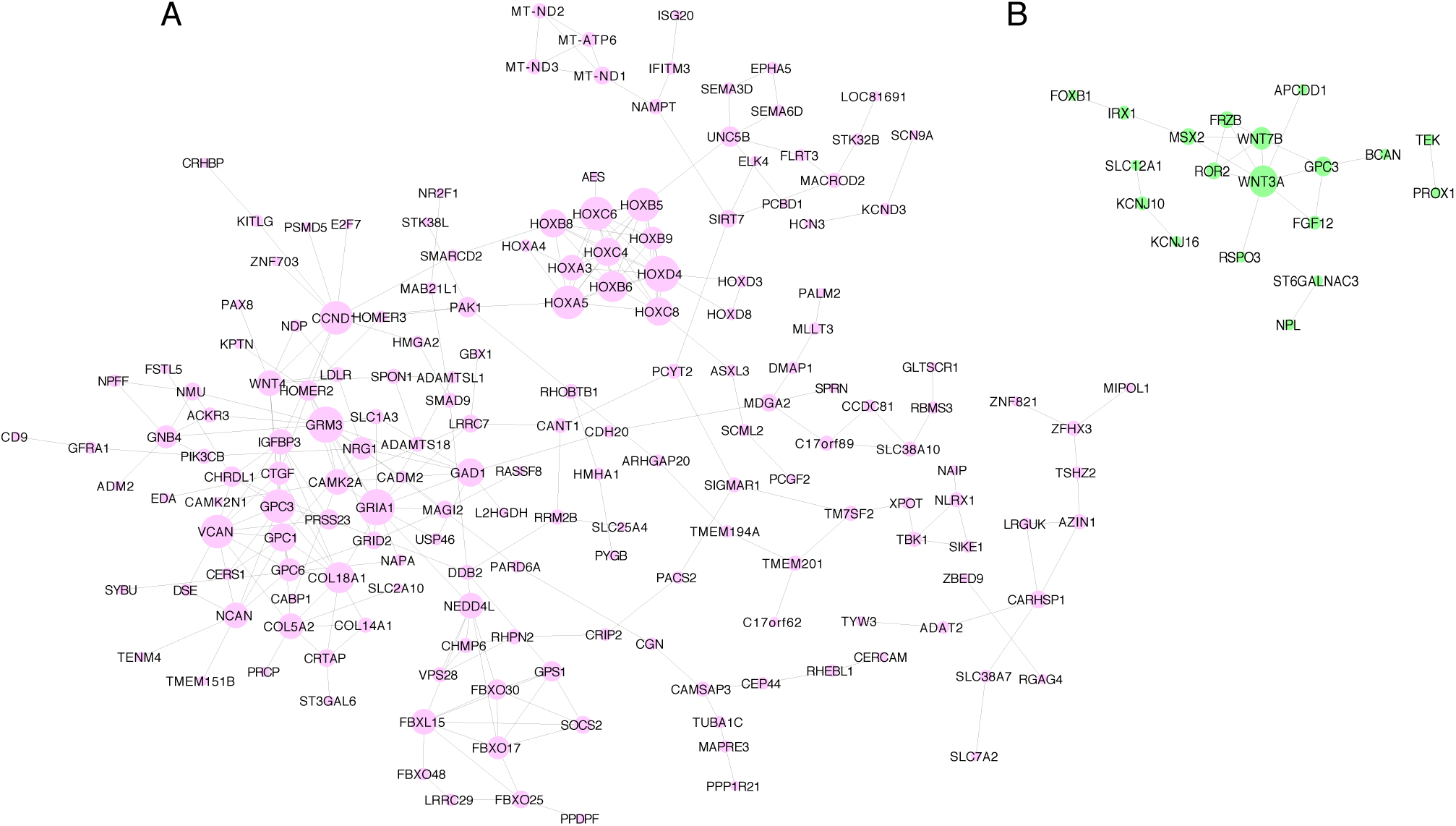
Protein-protein interaction network. Direct protein–protein interaction network of differentially expressed genes identified in (**A**) iNPCs and (**B**) iNeurons.

**Figure S8.**
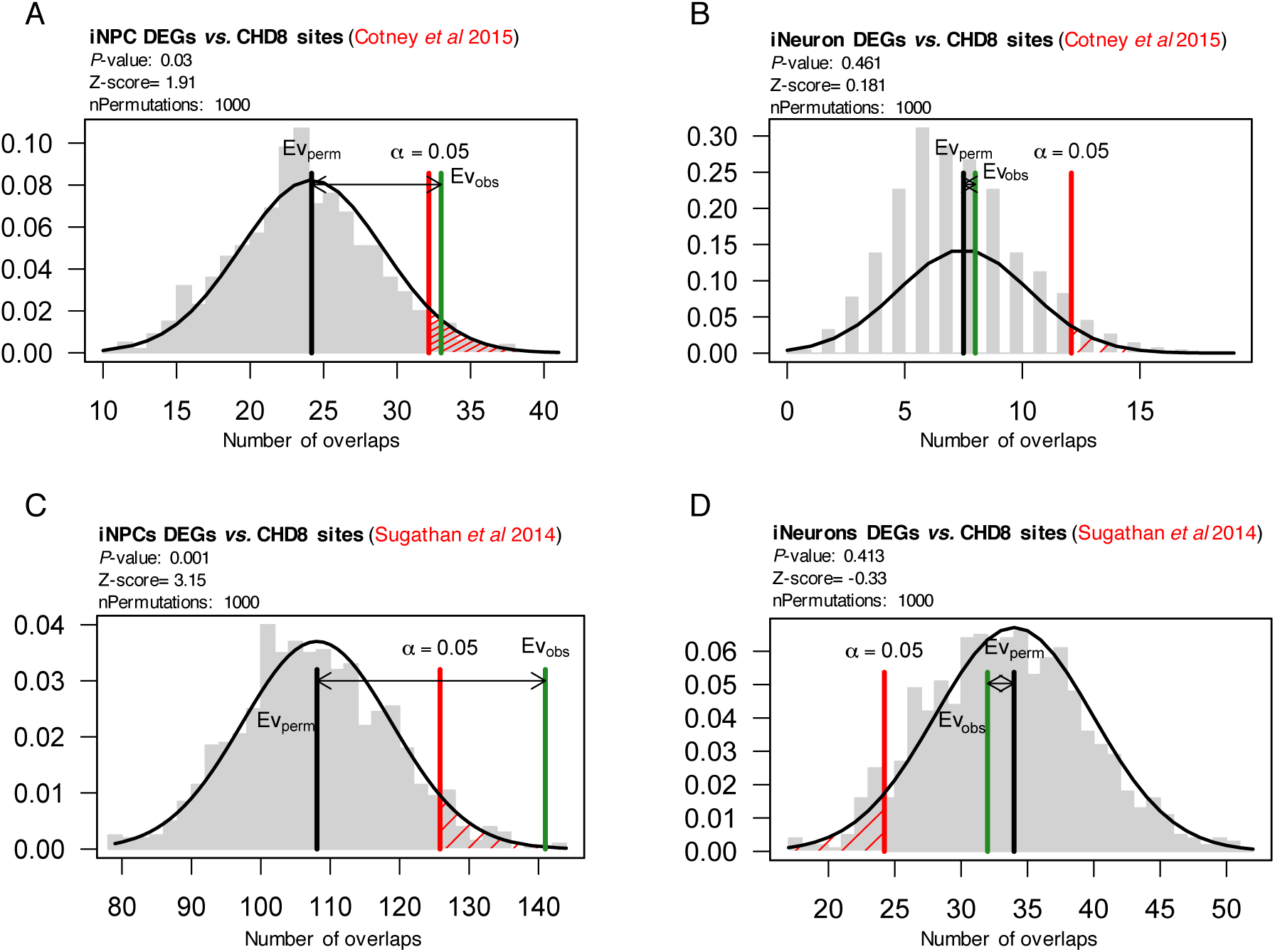
CHD8 enrichment analysis. Genomic coordinates for differentially expressed genes in iNPCs and iNeurons were assessed for enrichment for human brain specific CHD8 binding sites derived from (**A-B**) human mid-fetal brain tissue and (**C-D**) human neural progenitor cells (NPCs). The regioneR R package was used to test overlaps of genomic regions based on permutation sampling. We sampled random regions from the genome 1000 times, matching size and chromosomal distribution of the region set under study. By recomputing the overlap with *CHD8* binding sites in each permutation, statistical significance of the observed overlap was computed.

**Figure S9.**
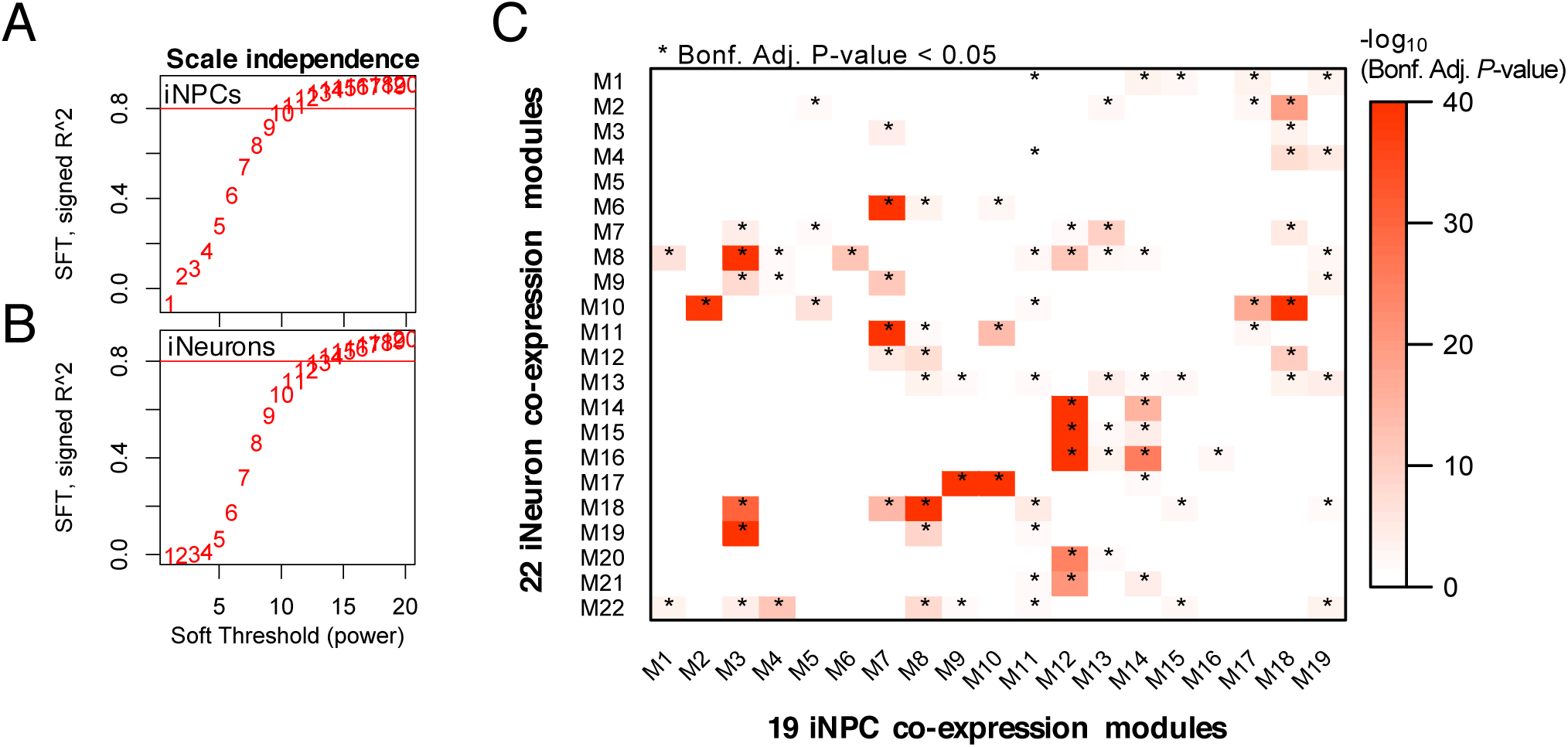
WGCNA module construction and overlap. The β-power defined for both (**A**) iNPC and (**B**) iNeuron samples in order to achieve scale free network topology for gene co-expression network construction. As a rule of thumb, β-power’s > 0.8 achieve scale free network topology, and a final β-power of 12 was used iNPC samples and a β-power of 14 for iNeuron samples. (**C**) Overlap analysis of co-expression modules defined based on iNPC and iNeuron samples. Significance of the overlap was tested using a one-sided Fisher’s exact test and corrected for multiple comparisons using Bonferroni procedure. Significant overlaps with adjusted *P*<0.05 are marked with an asterisks (*).

**Figure S10.**
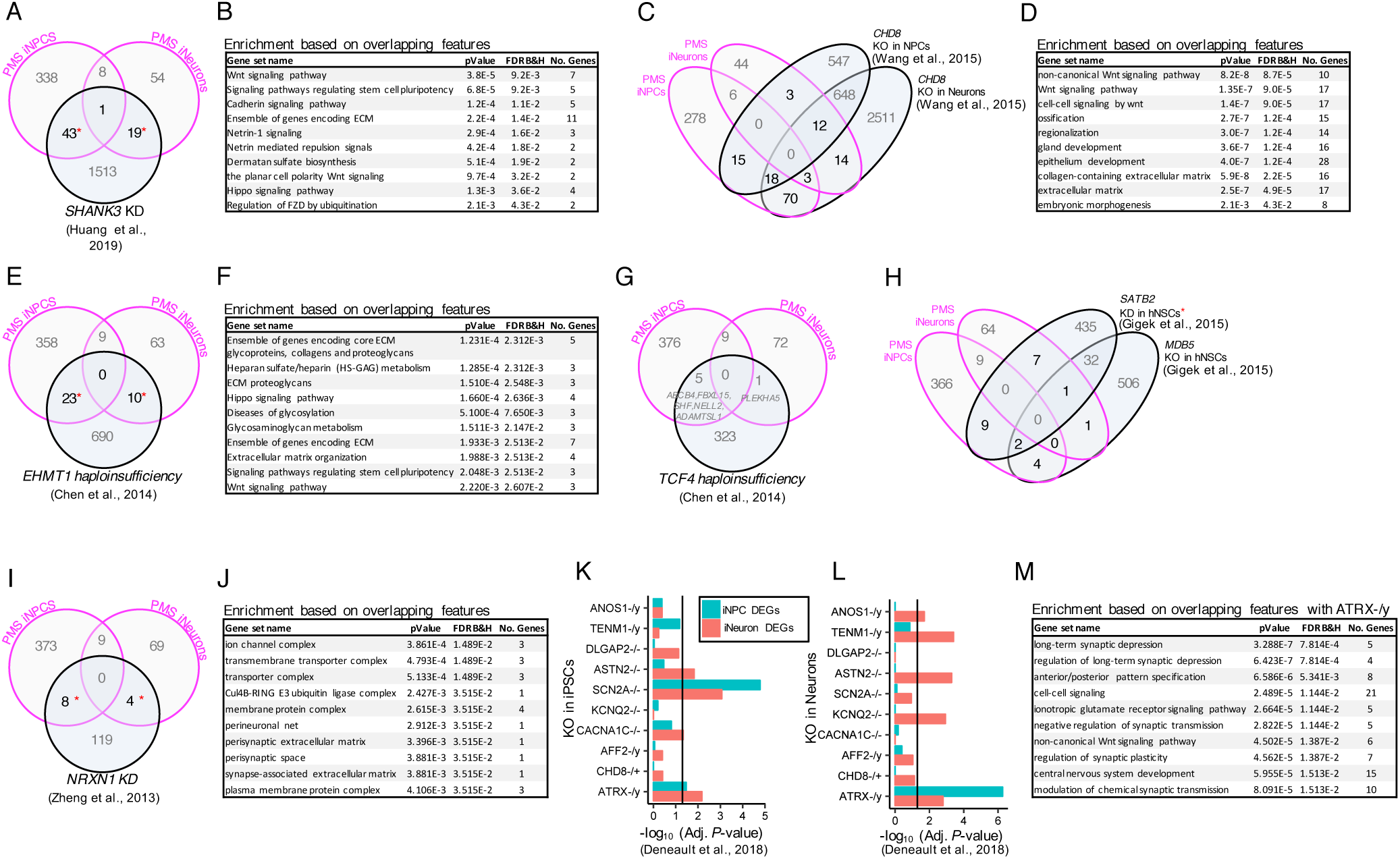
Overlap with other ASD iPSC transcriptome studies. Convergence of differentially expressed genes (FDR <5%) in the current study with other ASD iPSC transcriptome studies was assessed using a Fisher’s Exact Test (FET) and an estimated odds-ratio was computed in comparison to a genome-wide background set to 20,000. Significant overlaps are demarked with a red asterisks (*). All overlapping genes found in common with the currents study were subjected to ToppGene functional enrichment. Overlap of differentially expressed genes and functional annotation was performed using data from (**A-B**) Huang et al., 2019, (**C-D**) Wang et al., 2015, and (**E-G**) Chen et al., 2014, (**H**) Gigek et al., 2015, (**I-J**) Zheng et al., 2013, and (**K-M**) Deneault et al., 2018. Note that no functional enrichment was observed based on the overlap with (**G**) Chen et al., 2014 and (**H**) Gigek et al., 2015. To simply multiple overlaps, (**K-L**) the −log_10_ P-value (x-axis) is used to display the extent of significance based on gene expression perturbations associated with CRISPR/Cas9 knockout of 10 different ASD genes.

**Figure S11.**
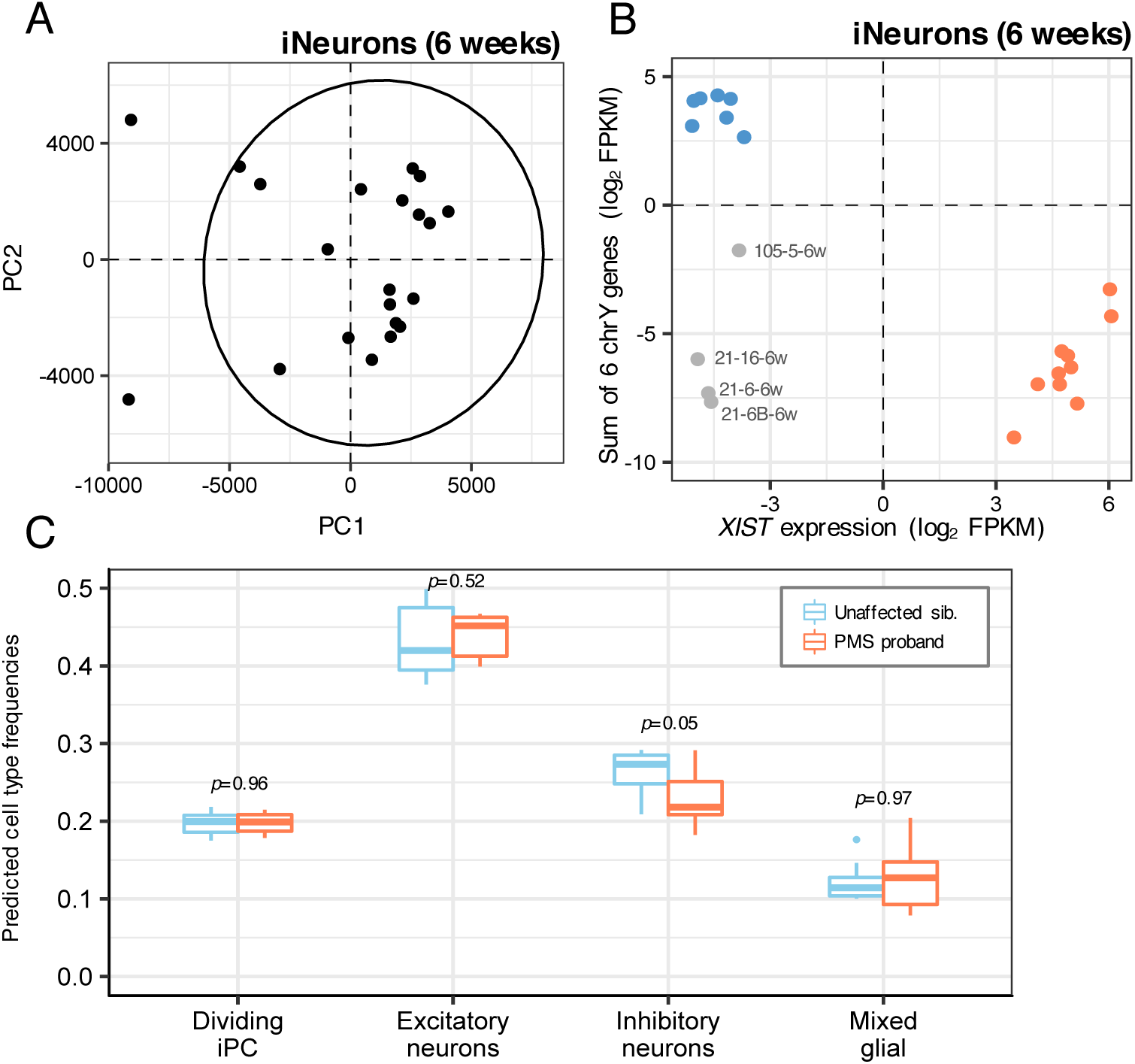
Data pre-processing using replication iNeuron samples. Principal component analyses were performed on RPKM values for (**A**) all replication set iNeuron samples at 6 weeks. Outliers beyond the 95% confidence intervals (black ellipse) were excluded from downstream analyses. (**B**) We also sought to identify samples that may have under-gone issues with X-inactivation and/or sample mislabeling by confirming that the reported biological sex is concordant with gene expression on chrX and chrY, which confirmed aberrant X-inactivation observed in iNPCs sharing the same clone and induction (*Supplemental Table 1*). Samples with intermediate expression profiles were excluded from further analysis. (**C**) Cibersort cell type deconvolution analysis of global gene expression profiles estimated cell frequencies (y-axis) for four major cell types (x-axis) using a reference panel of single-cell RNA-sequencing data from the human fetal cortex. The predicted cellular proportions were compared between PMS probands and unaffected siblings using a Wilcox rank-sum test.

## REFERENCES

1. Betancur C, Buxbaum JD: SHANK3 haploinsufficiency: a “common” but underdiagnosed highly penetrant monogenic cause of autism spectrum disorders. Mol Autism 2013, 4(1):17.

2. Leblond CS, Nava C, Polge A, Gauthier J, Huguet G, Lumbroso S, Giuliano F, Stordeur C, Depienne C, Mouzat K et al: Meta-analysis of SHANK Mutations in Autism Spectrum Disorders: a gradient of severity in cognitive impairments. PLoS Genet 2014, 10(9):e1004580.

3. Boccuto L, Lauri M, Sarasua SM, Skinner CD, Buccella D, Dwivedi A, Orteschi D, Collins JS, Zollino M, Visconti P et al: Prevalence of SHANK3 variants in patients with different subtypes of autism spectrum disorders. Eur J Hum Genet 2013, 21(3):310–316.

4. De Rubeis S, Siper PM, Durkin A, Weissman J, Muratet F, Halpern D, Trelles MDP, Frank Y, Lozano R, Wang AT et al: Delineation of the genetic and clinical spectrum of Phelan-McDermid syndrome caused by SHANK3 point mutations. Mol Autism 2018, 9:31.

5. Mitz AR, Philyaw TJ, Boccuto L, Shcheglovitov A, Sarasua SM, Kaufmann WE, Thurm A: Identification of 22q13 genes most likely to contribute to Phelan McDermid syndrome. Eur J Hum Genet 2018, 26(3):293–302.

6. Soorya L, Kolevzon A, Zweifach J, Lim T, Dobry Y, Schwartz L, Frank Y, Wang AT, Cai G, Parkhomenko E, et al: Prospective investigation of autism and genotype-phenotype correlations in 22q13 deletion syndrome and SHANK3 deficiency. Mol Autism 2013, 4(1):18.

7. Sarasua SM, Dwivedi A, Boccuto L, Chen CF, Sharp JL, Rollins JD, Collins JS, Rogers RC, Phelan K, DuPont BR: 22q13.2q13.32 genomic regions associated with severity of speech delay, developmental delay, and physical features in Phelan-McDermid syndrome. Genet Med 2014, 16(4):318–328.

8. Durand CM, Perroy J, Loll F, Perrais D, Fagni L, Bourgeron T, Montcouquiol M, Sans N: SHANK3 mutations identified in autism lead to modification of dendritic spine morphology via an actin-dependent mechanism. Mol Psychiatry 2012, 17(1):71–84.

9. Halbedl S, Schoen M, Feiler MS, Boeckers TM, Schmeisser MJ: Shank3 is localized in axons and presynaptic specializations of developing hippocampal neurons and involved in the modulation of NMDA receptor levels at axon terminals. J Neurochem 2016, 137(1):26–32.

10. Monteiro P, Feng G: SHANK proteins: roles at the synapse and in autism spectrum disorder. Nat Rev Neurosci 2017, 18(3):147–157.

11. Lee K, Vyas Y, Garner CC, Montgomery JM: Autism-associated Shank3 mutations alter mGluR expression and mGluR-dependent but not NMDA receptor-dependent long-term depression. Synapse 2019, 73(8):e22097.

12. Ponna SK, Ruskamo S, Myllykoski M, Keller C, Boeckers TM, Kursula P: Structural basis for PDZ domain interactions in the post-synaptic density scaffolding protein Shank3. J Neurochem 2018, 145(6):449–463.

13. Shi R, Redman P, Ghose D, Hwang H, Liu Y, Ren X, Ding LJ, Liu M, Jones KJ, Xu W: Shank Proteins Differentially Regulate Synaptic Transmission. eNeuro 2017, 4(6).

14. Verpelli C, Dvoretskova E, Vicidomini C, Rossi F, Chiappalone M, Schoen M, Di Stefano B, Mantegazza R, Broccoli V, Bockers TM et al: Importance of Shank3 protein in regulating metabotropic glutamate receptor 5 (mGluR5) expression and signaling at synapses. J Biol Chem 2011, 286(40):34839–34850.

15. Bey AL, Wang X, Yan H, Kim N, Passman RL, Yang Y, Cao X, Towers AJ, Hulbert SW, Duffney LJ et al: Brain region-specific disruption of Shank3 in mice reveals a dissociation for cortical and striatal circuits in autism-related behaviors. Transl Psychiatry 2018, 8(1):94.

16. Bozdagi O, Sakurai T, Papapetrou D, Wang X, Dickstein DL, Takahashi N, Kajiwara Y, Yang M, Katz AM, Scattoni ML et al: Haploinsufficiency of the autism-associated Shank3 gene leads to deficits in synaptic function, social interaction, and social communication. Mol Autism 2010, 1(1):15.

17. Dhamne SC, Silverman JL, Super CE, Lammers SHT, Hameed MQ, Modi ME, Copping NA, Pride MC, Smith DG, Rotenberg A et al: Replicable in vivo physiological and behavioral phenotypes of the Shank3B null mutant mouse model of autism. Mol Autism 2017, 8:26.

18. Drapeau E, Riad M, Kajiwara Y, Buxbaum JD: Behavioral Phenotyping of an Improved Mouse Model of Phelan-McDermid Syndrome with a Complete Deletion of the Shank3 Gene. eNeuro 2018, 5(3).

19. Ferhat AT, Halbedl S, Schmeisser MJ, Kas MJ, Bourgeron T, Ey E: Behavioural Phenotypes and Neural Circuit Dysfunctions in Mouse Models of Autism Spectrum Disorder. Adv Anat Embryol Cell Biol 2017, 224:85–101.

20. Jaramillo TC, Speed HE, Xuan Z, Reimers JM, Liu S, Powell CM: Altered Striatal Synaptic Function and Abnormal Behaviour in Shank3 Exon4-9 Deletion Mouse Model of Autism. Autism Res 2016, 9(3):350–375.

21. Kouser M, Speed HE, Dewey CM, Reimers JM, Widman AJ, Gupta N, Liu S, Jaramillo TC, Bangash M, Xiao B et al: Loss of predominant Shank3 isoforms results in hippocampus-dependent impairments in behavior and synaptic transmission. J Neurosci 2013, 33(47):18448–18468.

22. Lee J, Chung C, Ha S, Lee D, Kim DY, Kim H, Kim E: Shank3-mutant mice lacking exon 9 show altered excitation/inhibition balance, enhanced rearing, and spatial memory deficit. Front Cell Neurosci 2015, 9:94.

23. Peca J, Feliciano C, Ting JT, Wang W, Wells MF, Venkatraman TN, Lascola CD, Fu Z, Feng G: Shank3 mutant mice display autistic-like behaviours and striatal dysfunction. Nature 2011, 472(7344):437–442.

24. Speed HE, Kouser M, Xuan Z, Reimers JM, Ochoa CF, Gupta N, Liu S, Powell CM: Autism-Associated Insertion Mutation (InsG) of Shank3 Exon 21 Causes Impaired Synaptic Transmission and Behavioral Deficits. J Neurosci 2015, 35(26):9648–9665.

25. Wang X, Bey AL, Katz BM, Badea A, Kim N, David LK, Duffney LJ, Kumar S, Mague SD, Hulbert SW et al: Altered mGluR5-Homer scaffolds and corticostriatal connectivity in a Shank3 complete knockout model of autism. Nat Commun 2016, 7:11459.

26. Wang X, McCoy PA, Rodriguiz RM, Pan Y, Je HS, Roberts AC, Kim CJ, Berrios J, Colvin JS, Bousquet-Moore D et al: Synaptic dysfunction and abnormal behaviors in mice lacking major isoforms of Shank3. Hum Mol Genet 2011, 20(15):3093–3108.

27. Harony-Nicolas H, Kay M, du Hoffmann J, Klein ME, Bozdagi-Gunal O, Riad M, Daskalakis NP, Sonar S, Castillo PE, Hof PR, et al: Oxytocin improves behavioral and electrophysiological deficits in a novel Shank3-deficient rat. Elife 2017, 6.

28. Berg EL, Copping NA, Rivera JK, Pride MC, Careaga M, Bauman MD, Berman RF, Lein PJ, Harony-Nicolas H, Buxbaum JD et al: Developmental social communication deficits in the Shank3 rat model of phelan-mcdermid syndrome and autism spectrum disorder. Autism Res 2018, 11(4):587–601.

29. Mokhtari R, Lachman HM: Neurons Derived From Patient-Specific Induced Pluripotent Stem Cells: a Promising Strategy Towards Developing Novel Pharmacotherapies for Autism Spectrum Disorders. EBioMedicine 2016, 9:21–22.

30. Shen X, Yeung HT, Lai KO: Application of Human-Induced Pluripotent Stem Cells (hiPSCs) to Study Synaptopathy of Neurodevelopmental Disorders. Dev Neurobiol 2019, 79(1):20–35.

31. Shcheglovitov A, Shcheglovitova O, Yazawa M, Portmann T, Shu R, Sebastiano V, Krawisz A, Froehlich W, Bernstein JA, Hallmayer JF et al: SHANK3 and IGF1 restore synaptic deficits in neurons from 22q13 deletion syndrome patients. Nature 2013, 503(7475):267–271.

32. Kathuria A, Nowosiad P, Jagasia R, Aigner S, Taylor RD, Andreae LC, Gatford NJF, Lucchesi W, Srivastava DP, Price J: Stem cell-derived neurons from autistic individuals with SHANK3 mutation show morphogenetic abnormalities during early development. Mol Psychiatry 2018, 23(3):735–746.

33. Gouder L, Vitrac A, Goubran-Botros H, Danckaert A, Tinevez JY, Andre-Leroux G, Atanasova E, Lemiere N, Biton A, Leblond CS et al: Altered spinogenesis in iPSC-derived cortical neurons from patients with autism carrying de novo SHANK3 mutations. Sci Rep 2019, 9(1):94.

34. Darville H, Poulet A, Rodet-Amsellem F, Chatrousse L, Pernelle J, Boissart C, Heron D, Nava C, Perrier A, Jarrige M, et al: Human Pluripotent Stem Cell-derived Cortical Neurons for High Throughput Medication Screening in Autism: A Proof of Concept Study in SHANK3 Haploinsufficiency Syndrome. EBioMedicine 2016, 9:293–305.

35. Bidinosti M, Botta P, Kruttner S, Proenca CC, Stoehr N, Bernhard M, Fruh I, Mueller M, Bonenfant D, Voshol H et al: CLK2 inhibition ameliorates autistic features associated with SHANK3 deficiency. Science 2016, 351(6278):1199–1203.

36. Vicidomini C, Ponzoni L, Lim D, Schmeisser MJ, Reim D, Morello N, Orellana D, Tozzi A, Durante V, Scalmani P et al: Homer1b/c clustering is impaired in Phelan-McDermid Syndrome iPSCs derived neurons. Mol Psychiatry 2017, 22(5):637.

37. Yi F, Danko T, Botelho SC, Patzke C, Pak C, Wernig M, Sudhof TC: Autism-associated SHANK3 haploinsufficiency causes Ih channelopathy in human neurons. Science 2016, 352(6286):aaf2669.

38. Yang W, Mills JA, Sullivan S, Liu Y, French DL, Gadue P: iPSC Reprogramming from Human Peripheral Blood Using Sendai Virus Mediated Gene Transfer. In: StemBook. Cambridge (MA); 2008.

39. Bock C, Kiskinis E, Verstappen G, Gu H, Boulting G, Smith ZD, Ziller M, Croft GF, Amoroso MW, Oakley DH et al: Reference Maps of human ES and iPS cell variation enable high-throughput characterization of pluripotent cell lines. Cell 2011, 144(3):439–452.

40. Brennand KJ, Simone A, Jou J, Gelboin-Burkhart C, Tran N, Sangar S, Li Y, Mu Y, Chen G, Yu D et al: Modelling schizophrenia using human induced pluripotent stem cells. Nature 2011, 473(7346):221–225.

41. Bolger AM, Lohse M, Usadel B: Trimmomatic: a flexible trimmer for Illumina sequence data. Bioinformatics 2014, 30(15):2114–2120.

42. Dobin A, Davis CA, Schlesinger F, Drenkow J, Zaleski C, Jha S, Batut P, Chaisson M, Gingeras TR: STAR: ultrafast universal RNA-seq aligner. Bioinformatics 2013, 29(1):15–21.

43. Liao Y, Smyth GK, Shi W: featureCounts: an efficient general purpose program for assigning sequence reads to genomic features. Bioinformatics 2014, 30(7):923–930.

44. Hoffman GE, Hartley BJ, Flaherty E, Ladran I, Gochman P, Ruderfer DM, Stahl EA, Rapoport J, Sklar P, Brennand KJ: Transcriptional signatures of schizophrenia in hiPSC-derived NPCs and neurons are concordant with post-mortem adult brains. Nat Commun 2017, 8(1):2225.

45. Ritchie ME, Phipson B, Wu D, Hu Y, Law CW, Shi W, Smyth GK: limma powers differential expression analyses for RNA-sequencing and microarray studies. Nucleic Acids Res 2015, 43(7):e47.

46. Satterstrom FK, Kosmicki J, Wang J, Breen MS, De Rubeis S, An JY, Peng M, Collins R, Grove J, Klei L et al: Large-scale exome sequencing study implicates both developmental and functional changes in the neurobiology of autism. bioRxiv 2019, 484113(doi: https://doi.org/10.1101/484113).

47. Newman AM, Liu CL, Green MR, Gentles AJ, Feng W, Xu Y, Hoang CD, Diehn M, Alizadeh AA: Robust enumeration of cell subsets from tissue expression profiles. Nat Methods 2015, 12(5):453–457.

48. Nowakowski TJ, Bhaduri A, Pollen AA, Alvarado B, Mostajo-Radji MA, Di Lullo E, Haeussler M, Sandoval-Espinosa C, Liu SJ, Velmeshev D et al: Spatiotemporal gene expression trajectories reveal developmental hierarchies of the human cortex. Science 2017, 358(6368):1318–1323.

49. Hoffman GE, Schadt EE: variancePartition: interpreting drivers of variation in complex gene expression studies. BMC Bioinformatics 2016, 17(1):483.

50. Chen J, Bardes EE, Aronow BJ, Jegga AG: ToppGene Suite for gene list enrichment analysis and candidate gene prioritization. Nucleic Acids Res 2009, 37(Web Server issue):W305–311.

51. Szklarczyk D, Gable AL, Lyon D, Junge A, Wyder S, Huerta-Cepas J, Simonovic M, Doncheva NT, Morris JH, Bork P et al: STRING v11: protein-protein association networks with increased coverage, supporting functional discovery in genome-wide experimental datasets. Nucleic Acids Res 2019, 47(D1):D607–D613.

52. Shannon P, Markiel A, Ozier O, Baliga NS, Wang JT, Ramage D, Amin N, Schwikowski B, Ideker T: Cytoscape: a software environment for integrated models of biomolecular interaction networks. Genome Res 2003, 13(11):2498–2504.

53. Cotney J, Muhle RA, Sanders SJ, Liu L, Willsey AJ, Niu W, Liu W, Klei L, Lei J, Yin J et al: The autism-associated chromatin modifier CHD8 regulates other autism risk genes during human neurodevelopment. Nat Commun 2015, 6:6404.

54. Sugathan A, Biagioli M, Golzio C, Erdin S, Blumenthal I, Manavalan P, Ragavendran A, Brand H, Lucente D, Miles J et al: CHD8 regulates neurodevelopmental pathways associated with autism spectrum disorder in neural progenitors. Proc Natl Acad Sci U S A 2014, 111(42):E4468–4477.

55. Gel B, Diez-Villanueva A, Serra E, Buschbeck M, Peinado MA, Malinverni R: regioneR: an R/Bioconductor package for the association analysis of genomic regions based on permutation tests. Bioinformatics 2016, 32(2):289–291.

56. Langfelder P, Horvath S: WGCNA: an R package for weighted correlation network analysis. BMC Bioinformatics 2008, 9:559.

57. Betancur C: Etiological heterogeneity in autism spectrum disorders: more than 100 genetic and genomic disorders and still counting. Brain Res 2011, 1380:42–77.

58. De Rubeis S, He X, Goldberg AP, Poultney CS, Samocha K, Cicek AE, Kou Y, Liu L, Fromer M, Walker S et al: Synaptic, transcriptional and chromatin genes disrupted in autism. Nature 2014, 515(7526):209–215.

59. Gilman SR, Iossifov I, Levy D, Ronemus M, Wigler M, Vitkup D: Rare de novo variants associated with autism implicate a large functional network of genes involved in formation and function of synapses. Neuron 2011, 70(5):898–907.

60. Iossifov I, O’Roak BJ, Sanders SJ, Ronemus M, Krumm N, Levy D, Stessman HA, Witherspoon KT, Vives L, Patterson KE et al: The contribution of de novo coding mutations to autism spectrum disorder. Nature 2014, 515(7526):216–221.

61. Parikshak NN, Luo R, Zhang A, Won H, Lowe JK, Chandran V, Horvath S, Geschwind DH: Integrative functional genomic analyses implicate specific molecular pathways and circuits in autism. Cell 2013, 155(5):1008–1021.

62. Pinto D, Delaby E, Merico D, Barbosa M, Merikangas A, Klei L, Thiruvahindrapuram B, Xu X, Ziman R, Wang Z et al: Convergence of genes and cellular pathways dysregulated in autism spectrum disorders. Am J Hum Genet 2014, 94(5):677–694.

63. Wright CF, Fitzgerald TW, Jones WD, Clayton S, McRae JF, van Kogelenberg M, King DA, Ambridge K, Barrett DM, Bayzetinova T et al: Genetic diagnosis of developmental disorders in the DDD study: a scalable analysis of genome-wide research data. Lancet 2015, 385(9975):1305–1314.

64. Darnell JC, Van Driesche SJ, Zhang C, Hung KY, Mele A, Fraser CE, Stone EF, Chen C, Fak JJ, Chi SW et al: FMRP stalls ribosomal translocation on mRNAs linked to synaptic function and autism. Cell 2011, 146(2):247–261.

65. Huang G, Chen S, Chen X, Zheng J, Xu Z, Doostparast Torshizi A, Gong S, Chen Q, Ma X, Yu J et al: Uncovering the Functional Link Between SHANK3 Deletions and Deficiency in Neurodevelopment Using iPSC-Derived Human Neurons. Front Neuroanat 2019, 13:23.

66. Wang P, Lin M, Pedrosa E, Hrabovsky A, Zhang Z, Guo W, Lachman HM, Zheng D: CRISPR/Cas9-mediated heterozygous knockout of the autism gene CHD8 and characterization of its transcriptional networks in neurodevelopment. Mol Autism 2015, 6:55.

67. Chen ES, Gigek CO, Rosenfeld JA, Diallo AB, Maussion G, Chen GG, Vaillancourt K, Lopez JP, Crapper L, Poujol R et al: Molecular convergence of neurodevelopmental disorders. Am J Hum Genet 2014, 95(5):490–508.

68. Gigek CO, Chen ES, Ota VK, Maussion G, Peng H, Vaillancourt K, Diallo AB, Lopez JP, Crapper L, Vasuta C et al: A molecular model for neurodevelopmental disorders. Transl Psychiatry 2015, 5:e565.

69. Zeng L, Zhang P, Shi L, Yamamoto V, Lu W, Wang K: Functional impacts of NRXN1 knockdown on neurodevelopment in stem cell models. PLoS One 2013, 8(3):e59685.

70. Deneault E, White SH, Rodrigues DC, Ross PJ, Faheem M, Zaslavsky K, Wang Z, Alexandrova R, Pellecchia G, Wei W et al: Complete Disruption of Autism-Susceptibility Genes by Gene Editing Predominantly Reduces Functional Connectivity of Isogenic Human Neurons. Stem Cell Reports 2019, 12(2):427–429.

71. Ha HTT, Leal-Ortiz S, Lalwani K, Kiyonaka S, Hamachi I, Mysore SP, Montgomery JM, Garner CC, Huguenard JR, Kim SA: Shank and Zinc Mediate an AMPA Receptor Subunit Switch in Developing Neurons. Front Mol Neurosci 2018, 11:405.

72. Wu S, Gan G, Zhang Z, Sun J, Wang Q, Gao Z, Li M, Jin S, Huang J, Thomas U et al: A Presynaptic Function of Shank Protein in Drosophila. J Neurosci 2017, 37(48):11592–11604.

73. Kozol RA, Cukier HN, Zou B, Mayo V, De Rubeis S, Cai G, Griswold AJ, Whitehead PL, Haines JL, Gilbert JR et al: Two knockdown models of the autism genes SYNGAP1 and SHANK3 in zebrafish produce similar behavioral phenotypes associated with embryonic disruptions of brain morphogenesis. Hum Mol Genet 2015, 24(14):4006–4023.

74. Arons MH, Thynne CJ, Grabrucker AM, Li D, Schoen M, Cheyne JE, Boeckers TM, Montgomery JM, Garner CC: Autism-associated mutations in ProSAP2/Shank3 impair synaptic transmission and neurexin-neuroligin-mediated transsynaptic signaling. J Neurosci 2012, 32(43):14966–14978.

75. Betancur C, Sakurai T, Buxbaum JD: The emerging role of synaptic cell-adhesion pathways in the pathogenesis of autism spectrum disorders. Trends Neurosci 2009, 32(7):402–412.

76. Sudhof TC: Neuroligins and neurexins link synaptic function to cognitive disease. Nature 2008, 455(7215):903–911.

77. Meyer G, Varoqueaux F, Neeb A, Oschlies M, Brose N: The complexity of PDZ domain-mediated interactions at glutamatergic synapses: a case study on neuroligin. Neuropharmacology 2004, 47(5):724–733.

78. Budreck EC, Kwon OB, Jung JH, Baudouin S, Thommen A, Kim HS, Fukazawa Y, Harada H, Tabuchi K, Shigemoto R et al: Neuroligin-1 controls synaptic abundance of NMDA-type glutamate receptors through extracellular coupling. Proc Natl Acad Sci U S A 2013, 110(2):725–730.

79. Harris KP, Akbergenova Y, Cho RW, Baas-Thomas MS, Littleton JT: Shank Modulates Postsynaptic Wnt Signaling to Regulate Synaptic Development. J Neurosci 2016, 36(21):5820–5832.

80. Quesnel-Vallieres M, Weatheritt RJ, Cordes SP, Blencowe BJ: Autism spectrum disorder: insights into convergent mechanisms from transcriptomics. Nat Rev Genet 2019, 20(1):51–63.

81. Kalkman HO: A review of the evidence for the canonical Wnt pathway in autism spectrum disorders. Mol Autism 2012, 3(1):10.

82. Platt RJ, Zhou Y, Slaymaker IM, Shetty AS, Weisbach NR, Kim JA, Sharma J, Desai M, Sood S, Kempton HR et al: Chd8 Mutation Leads to Autistic-like Behaviors and Impaired Striatal Circuits. Cell Rep 2017, 19(2):335–350.

83. Chenn A, Walsh CA: Regulation of cerebral cortical size by control of cell cycle exit in neural precursors. Science 2002, 297(5580):365–369.

84. Dong F, Jiang J, McSweeney C, Zou D, Liu L, Mao Y: Deletion of CTNNB1 in inhibitory circuitry contributes to autism-associated behavioral defects. Hum Mol Genet 2016, 25(13):2738–2751.

85. Krumm N, O’Roak BJ, Shendure J, Eichler EE: A de novo convergence of autism genetics and molecular neuroscience. Trends Neurosci 2014, 37(2):95–105.

86. Frazier TW, Embacher R, Tilot AK, Koenig K, Mester J, Eng C: Molecular and phenotypic abnormalities in individuals with germline heterozygous PTEN mutations and autism. Mol Psychiatry 2015, 20(9):1132–1138.

87. Spinelli L, Black FM, Berg JN, Eickholt BJ, Leslie NR: Functionally distinct groups of inherited PTEN mutations in autism and tumour syndromes. J Med Genet 2015, 52(2):128–134.

88. Chen Y, Huang WC, Sejourne J, Clipperton-Allen AE, Page DT: Pten Mutations Alter Brain Growth Trajectory and Allocation of Cell Types through Elevated beta-Catenin Signaling. J Neurosci 2015, 35(28):10252–10267.

89. Kwan V, Unda BK, Singh KK: Wnt signaling networks in autism spectrum disorder and intellectual disability. J Neurodev Disord 2016, 8:45.

90. Snijders Blok L, Madsen E, Juusola J, Gilissen C, Baralle D, Reijnders MR, Venselaar H, Helsmoortel C, Cho MT, Hoischen A et al: Mutations in DDX3X Are a Common Cause of Unexplained Intellectual Disability with Gender-Specific Effects on Wnt Signaling. Am J Hum Genet 2015, 97(2):343–352.

91. Gao C, Chen YG: Dishevelled: The hub of Wnt signaling. Cell Signal 2010, 22(5):717–727.

92. Mines MA, Jope RS: Brain region differences in regulation of Akt and GSK3 by chronic stimulant administration in mice. Cell Signal 2012, 24(7):1398–1405.

93. Zhou WJ, Xu N, Kong L, Sun SC, Xu XF, Jia MZ, Wang Y, Chen ZY: The antidepressant roles of Wnt2 and Wnt3 in stress-induced depression-like behaviors. Transl Psychiatry 2016, 6(9):e892.

94. Roh MS, Seo MS, Kim Y, Kim SH, Jeon WJ, Ahn YM, Kang UG, Juhnn YS, Kim YS: Haloperidol and clozapine differentially regulate signals upstream of glycogen synthase kinase 3 in the rat frontal cortex. Exp Mol Med 2007, 39(3):353–360.

95. Launay JM, Mouillet-Richard S, Baudry A, Pietri M, Kellermann O: Raphe-mediated signals control the hippocampal response to SRI antidepressants via miR-16. Transl Psychiatry 2011, 1:e56.

96. Sutton LP, Honardoust D, Mouyal J, Rajakumar N, Rushlow WJ: Activation of the canonical Wnt pathway by the antipsychotics haloperidol and clozapine involves dishevelled-3. J Neurochem 2007, 102(1):153–169.

